# In-depth Analysis of the Sirtuin 5-regulated Mouse Brain Acylome using Library-free Data-Independent Acquisitions

**DOI:** 10.1101/2022.08.06.503046

**Authors:** Joanna Bons, Jacob Rose, Ran Zhang, Jordan B. Burton, Christopher Carrico, Eric Verdin, Birgit Schilling

**Affiliations:** Buck Institute for Research on Aging, 8001 Redwood Boulevard, Novato, California 94945, USA

**Author notes:** Corresponding author: Birgit Schilling, Buck Institute for Research on Aging, 8001 Redwood Boulevard, Novato, California 94945, USA.

**Keywords:** Acylome, Brain, Data-independent acquisition, Post-translational modifications, Sirtuin 5

## Abstract

Post-translational modifications (PTMs) dynamically regulate proteins and biological pathways, typically through the combined effects of multiple PTMs. Lysine residues are targeted for various PTMs, including malonylation and succinylation. However, PTMs offer specific challenges to mass spectrometry-based proteomics during data acquisition and processing. Thus, novel and innovative workflows using data-independent acquisition (DIA) ensure confident PTM identification, precise site localization, and accurate and robust label-free quantification. In this study, we present a powerful approach that combines antibody-based enrichment with comprehensive DIA acquisitions and spectral library-free data processing using directDIA (Spectronaut). Identical DIA data can be used to generate spectral libraries and comprehensively identify and quantify PTMs, reducing the amount of enriched sample and acquisition time needed, while offering a fully automated workflow. We analyzed brains from wild-type and Sirtuin 5 (SIRT5)-knock-out mice, and discovered and quantified 466 malonylated and 2,211 succinylated peptides. SIRT5 regulation remodeled the acylomes by targeting 171 malonylated and 640 succinylated sites. Affected pathways included carbohydrate and lipid metabolisms, synaptic vesicle cycle, and neurodegenerative diseases. We found 48 common SIRT5-regulated malonylation and succinylation sites, suggesting potential PTM crosstalk. This innovative and efficient workflow offers deeper insights into the mouse brain lysine malonylome and succinylome.

**Statement of significance of the study:** Post-translational modifications (PTMs) are key regulators of protein structure, functions, and interactions. A great variety of PTMs have been discovered, including lysine acylation, such as acetylation, malonylation, and succinylation. Lysine acylation is understudied, particularly in the brain, and analysis by mass spectrometry-based proteomics faces significant challenges. In this study, we present a robust and efficient workflow to investigate proteome-wide PTM remodeling combining affinity PTM enrichment and a novel spectral library-free data-independent acquisition (DIA) approach. The strength of label-free DIA becomes evident with the collection of comprehensive information by tandem mass spectrometry for all detectable precursor ions of all biological samples, and the highly accurate quantitative information that can subsequently be retrieved with time-efficient and straightforward library-free strategies. More importantly, this enables confident identification of PTM sites and differentiation of PTM isomers. We applied this workflow to decipher the malonylome and succinylome remodeling and cross-talk in brains from wild-type and Sirt5^(-/-)^ mice, taking advantage of the demalonylase and desuccinylase activities of SIRT5, a nicotinamide adenine dinucleotide (NAD^+^)-dependent sirtuin. Interestingly, 10 malonylated proteins and 33 succinylated proteins targeted by SIRT5 are involved in the Parkinson’s disease pathway, including subunit beta of the calcium/calmodulin-dependent protein kinase type II (Camk2b) and protein DJ-1 (Park7).

## 1. Introduction

Protein post-translational modifications (PTMs) are important and dynamic regulators of protein structure, functions and interactions, that are involved in numerous biological processes including signaling, transcriptional regulation, protein trafficking and metabolism [1]. Hundreds of different PTMs have been identified [2-4], that typically modify various different amino acid residues. Acylation is a common covalent modification, and histone lysine acetylation was initially discovered to regulate transcription [5-8]. Over the last decade, significant improvements in mass spectrometry-based proteomics [4] and the development of highly specific antibodies for PTM enrichments enabled in-depth analysis and quantification of a multitude of PTMs [3, 9].

Initially, Zhao *et al*. developed an antibody against acetylated proteins [10], that opened the path to large acetylome and other acylomes studies [11-13], when combining the PTM enrichments with liquid chromatography-tandem mass spectrometry (LC-MS/MS). These reversible acyl modifications result from donation of a malonyl or succinyl group onto the ε-amino group of lysine sidechains either enzymatically by dedicated acyltransferases [14-16] or in non-enzymatic reactions [17-20]. These added negatively-charged carboxyl moieties change the net charge of lysine residues from a positive (+1) to a negative (−1) charge at physiological pH, which alter the physical and chemical properties of the proteins and impact their structures and interactions. Dysregulation of protein acylation is involved in cancer metastasis, diabetes and neurodegenerative diseases [16, 21].

Specific lysine deacylases catalyze the removal of the acylation groups, and include members of the sirtuin family [16, 22]. Sirtuins are a class of nicotinamide adenine dinucleotide (NAD^+^)-dependent deacylases, with seven members in mammals (SIRT1-7) that differ in subcellular localization, activity and substrate specificity [21]. Sirtuin 5 (SIRT5) localizes in mitochondria, cytoplasm, nucleus [23, 24] and peroxisomes [25]. SIRT5 presents high levels of enzymatic activities, such as desuccinylase [11, 12, 24], demalonylase [11, 12, 26, 27] and deglutarylase [28], but low acetylase activity [29]. SIRT5 plays a role in metabolism homeostasis by regulating glycolysis, the TCA cycle, the electron transport chain and urea cycle, and by promoting fatty acid β-oxidation, ketone body production and ROS detoxification [30]. SIRT5 acts also as a crucial cardioprotector, neuroprotector and cancer regulator.

SIRT5-regulation of protein PTMs is important, but remodeling of the malonylome and succinylome in general is, however, still understudied. Data-dependent acquisition (DDA) has been commonly used, but the semi-stochastic nature poses particular difficulties for PTM analysis, reproducible measurements of low-abundant modified peptides throughout the sample cohort, and single amino acid resolution for PTM site localization. Moreover, multiplexing isobaric-labeling strategies, such as isobaric tags for relative and absolute quantitation [31] or tandem mass tag [32] have challenges, including ratio compression leading to reduced accuracy [33]. This was initially resolved by the acquisition of MS3 scans [34], but at the expense of higher duty cycle, lower identification results and dedicated MS platforms [35, 36].

In this context, label-free DIA using high-resolution, accurate-mass mass spectrometers and most recent DIA strategies represent powerful alternatives for simultaneous PTM identification, quantification and site localization for acylation sites as demonstrated by our group [37, 38] and for other PTMs, such as phosphorylation [39, 40] and ubiquitination [41, 42]. DIA relies on the co-isolation and co-fragmentation of all precursor ions contained in segment-like m/z windows, and all fragment ions are then analyzed together [43]. The instrument cycles throughout the defined mass ranges to obtain comprehensive and systematic measurements of all detectable precursor ions contained in the samples. DIA thus provides a deep and robust profiling of peptides, with highly accurate quantification [44, 45]. Dedicated and sophisticated workflows were developed to analyze the multiplexed DIA MS/MS spectra [46, 47], commonly using an experimentally generated spectral library, which requires additional acquisition time and sample material to be generated.

Here, we present an innovative and efficient label-free quantitative workflow for acylome profiling combining antibody-based PTM peptide enrichment, comprehensive DIA acquisitions, and library-free DIA data processing referred to as directDIA in Spectronaut. This strategy uses the same DIA data to generate a spectral library, then to accurately quantify and precisely localize acylation sites in a fully automated fashion [39]. We applied this workflow to investigate and decipher the dynamic remodeling of lysine malonylome and succinylome in whole brains of wild-type (WT) and Sirt5^(-/-)^ mouse. We also explored the cross-talk between these two acylations, highlighting potential PTM targets in neurodegenerative diseases.

## 2. Materials and Methods

### 2.1. Mouse Brains

The animal studies were performed according to protocols approved by the Institutional Animal Care and Use Committee. SIRT5 knock-out (KO) mice were obtained from the Jackson Laboratory (Strain #012757). Mice were housed (12-h light/dark cycle, 22°C) and given unrestricted access to water. Brain tissues were collected from 18-month-old females, after 6 hours of refeeding and 24 hours of fasting.

### 2.2. Protein Digestion and Desalting

Mouse brain tissues were collected from two different conditions with four biological replicates each: i) WT (n=4), ii) SIRT5^(-/-)^ (SIRT5-KO, n=4). Frozen brains were homogenized in lysis buffer containing 8 M urea, 50 mM Tris, pH 7.5, 1x HALT protease inhibitor cocktail (Thermo Fisher Scientific, Waltham, MA), 150 mM NaCl, 5 μM trichostatin A and 5 mM nicotinamide, and homogenized for two cycles with a Bead Beater TissueLyser II (Qiagen, Germantown, MD) at 25 Hz for 3 min each. Lysates were clarified by spinning at 16,500 x *g* for 15 min at 4°C, and the supernatant containing the soluble proteins was collected. After protein concentration determination, 3 and 5 mg of protein from each sample were aliquoted for malonyl and succinyl peptide analysis, respectively. Proteins were reduced and alkylated before digesting overnight with a solution of sequencing-grade trypsin in 50 mM triethylammonium bicarbonate at a 1:50 (wt:wt) enzyme:protein ratio at 37°C. This reaction was quenched with 1% formic acid (FA), and the samples were clarified by centrifugation at 2,000 x *g* for 10 min at room temperature and desalted. Then 100 μg of each peptide elution were aliquoted for analysis of protein-level changes, and all desalted samples were vacuum dried. Then 100 µg of whole-lysate aliquots were re-suspended in 0.2% FA in water at a final concentration of 1 µg/µL and stored for MS analysis. The remaining 4.9 mg digests (for malonyl enrichment) and 2.9 mg digests (for succinyl enrichment) were re-suspended in 1.4 mL of immunoaffinity purification buffer (Cell Signaling Technology, Danvers, MA) containing 50 mM 4-morpholinepropanesulfonic acid (MOPS)/sodium hydroxide, pH 7.2, 10 mM disodium phosphate and 50 mM sodium chloride for PTM enrichment. Peptides were enriched for malonylation with anti-malonyl antibody conjugated to agarose beads from the Malonyl-Lysine Motif Kit (Kit #93872), and for succinylation with anti-succinyl antibody conjugated to agarose beads from the Succinyl-Lysine Motif Kit (Kit #13764; both from Cell Signaling Technology, Danvers, MA). This process was performed according to the manufacturer protocol; however, each sample was incubated in half the recommended volume of washed beads. Peptides were eluted from the antibody-bead conjugates with 0.1% trifluoroacetic acid in water and desalted. Samples were vacuum dried and re-suspended in 0.2% FA in water, before spiking indexed retention time standard peptides (Biognosys, Schlieren, Switzerland) [48] according to manufacturer’s instructions.

### 2.3. Mass Spectrometric Analysis

LC-MS/MS analyses were performed on a Dionex UltiMate 3000 system coupled to an Orbitrap Eclipse Tribrid mass spectrometer (both from Thermo Fisher Scientific, San Jose, CA). Proteolytic peptides (200 ng for the protein level analysis, and 4 µL for the PTM level analysis) were loaded onto an Acclaim PepMap 100 C18 trap column (0.1 × 20 mm, 5 µm particle size) for 5 min at 5 µL/min with 100% solvent A, and eluted on an Acclaim PepMap 100 C18 analytical column (75 µm x 50 cm, 3 µm particle size; both from Thermo Fisher Scientific) at 300 nL/min with the following gradient of solvent B (98% acetonitrile, 0.1% FA in H_2_O): 2% for 5 min, linear from 2% to 20% in 125 min, linear from 20% to 32% in 40 min, and up to 80% in 1 min, with a total gradient length of 210 min.

All samples, for protein and PTM level analyses, were acquired in DIA mode. Full MS spectra were collected at 120,000 resolution, and MS2 spectra at 30,000 resolution. The isolation scheme consisted in 26 variable windows covering the 350–1,650 m/z range with an overlap of 1 m/z (**Table S1**) [49]. Details about LC-MS/MS setups are given in **Supporting Information**. A detailed step-by-step procedure for building the DIA method and subsequent data processing can be found in **Appendix 1**.

### 2.4. Data Analysis with directDIA (Spectronaut)

DIA data was processed in Spectronaut (version 14.10.201222.47784) using directDIA for the protein level and PTM-enriched samples. Data was searched against the *Mus musculus* proteome with 58,430 protein entries (UniProtKB-TrEMBL), accessed on 01/31/2018. Trypsin/P was set as digestion enzyme and two missed cleavages were allowed. Cysteine carbamidomethylation was set as fixed modification, and methionine oxidation and protein N-terminus acetylation as variable modifications. Data extraction parameters were set as dynamic. Identification was performed using 1% precursor and protein q-value. For the protein lysate samples, quantification was based on the extracted ion chromatograms (XICs) of 3–6 MS2 fragment ions, local normalization was applied, and indexed retention time (iRT) profiling was selected. For the malonyl-enriched samples, lysine malonylation was additionally set as variable modification, and six missed cleavages were allowed. For the succinyl-enriched samples, lysine succinylation was additionally defined as variable modification, and four missed cleavages were allowed. PTM localization was selected with a probability cutoff of 0.75. Quantification was based on the XICs of 3–6 MS2 fragment ions, specifically b- and y-ions, without normalization, as well as data filtering with q-value sparse. Grouping and quantitation of PTM peptides were accomplished using the following criteria: minor grouping by modified sequence and minor group quantity by mean precursor quantity. A detailed step-by-step procedure for processing the DIA data, protein-level and PTM-level analyses, can be found in **Appendix 1**.

Differential expression analysis was performed using a paired t-test, and p-values were corrected for multiple testing [50]. For whole-lysate (protein-level) analysis, protein groups are required with at least two unique peptides. For determining differential changes, a q-value < 0.01 and absolute Log_2_(fold-change) > 0.58 was required at the protein level, and a q-value < 0.05 and absolute Log_2_(fold-change) > 0.58 at the PTM level.

### 2.5. Data analysis with Skyline

To increase the confidence of the malonyl peptide analysis, malonylated peptides identified using the described analysis were further manually investigated in Skyline [51]. To do so, a spectral library was first built in Spectronaut from the malonyl peptide DIA acquisitions using the Biognosys (BGS) default settings, except that lysine malonylation was added as variable modification and four missed cleavages were allowed, and contained 13,304 modified peptides and 3,112 protein groups (**Table S2**). The library was then imported into Skyline (Skyline-daily, version 21.2.1.377), and 468 malonylated peptides from 242 proteins and present in the library were finally targeted. At least three product ions (mono- and doubly-charged y- and b-type ions) from ion-2 to last ion-1 were extracted. All matching scans were used. Chromatographic peaks were manually checked to remove interfered transitions and correct peak boundaries. At least three transitions were kept per precursor ion. Peptide area was obtained by summing the corresponding transition peak areas.

Statistical analysis was performed in Skyline, and malonylated peptides with p < 0.05 and absolute fold-change > 1.5 were considered as significantly altered.

### 2.6. Clustering Analysis

Partial least squares-discriminant analysis of the proteomics data was performed using the package mixOmics [52] in R (version 4.0.2; RStudio, version 1.3.1093).

### 2.7. Enrichment Analysis

An over-representation analysis was performed using Consensus Path DB-mouse (Release MM11, 14.10.2021) [53, 54] to determine which gene ontology (GO) terms were significantly enriched. GO terms identified from the over-representation analysis were subjected to the following filters: q-value < 0.01 and term level > 3. Dot plots were generated using the ggplot2 package [55] in R (version 4.0.5; RStudio, version 1.4.1106). Kyoto Encyclopedia of Genes and Genomes (KEGG) pathway enrichment analysis was performed using the ClueGO package [56] (version 2.5.8), in Cytoscape [57] (version 3.8.2). Default settings were applied, except that two-sided hypergeometric test Bonferroni-adjusted p-value threshold was set to 0.01. Kappa score threshold was let at 0.4 for drawing pathway-connecting edges. Pathways with the same color indicate at least 50% similarity in genes/term.

### 2.8. iceLogo Heatmap

Heatmaps of sequences of 15 amino acids centered around the modified lysine residues were generated using the iceLogo tool [58] to determine the frequency of every amino acid residue around the modified lysine residue. The *M. musculus* precompiled Swiss-Prot composition was chosen, and the start position was set to 0. The p-value was set at 0.05, and significantly up- and downregulated residues were colored in shades of green and red, respectively.

### 2.9. Subcellular Localization

Subcellular localization was determined using Cytoscape [57] (version 3.8.2) and stringAPP [59] (version 1.7.0), by applying default settings, except that species was defined as *M. musculus*. Only compartments with scores above 4.5 were considered.

## 3. Results and Discussion

### 3.1. DIA-PTM workflow

To investigate malonylome and succinylome remodeling in mouse brain, an efficient and innovative workflow combining post-translationally modified peptide enrichment, comprehensive data-independent acquisitions (DIA) and spectral library-free DIA data analysis was developed, and is displayed on **Figure 1**. Four biological replicates of SIRT5-KO and four replicates of WT mouse brain were lysed using a urea-based buffer, proteins were digested in solution with trypsin, and proteolytic peptides were enriched for modified peptides, *i*.*e*. malonylated and succinylated peptides, as adapted from Holtz *et al*. [60] and Xie *et al*. [61], except that PTMScan malonyl-lysine or succinyl-lysine antibody beads (Cell Signaling Technology) were used.

**FIGURE 1.**
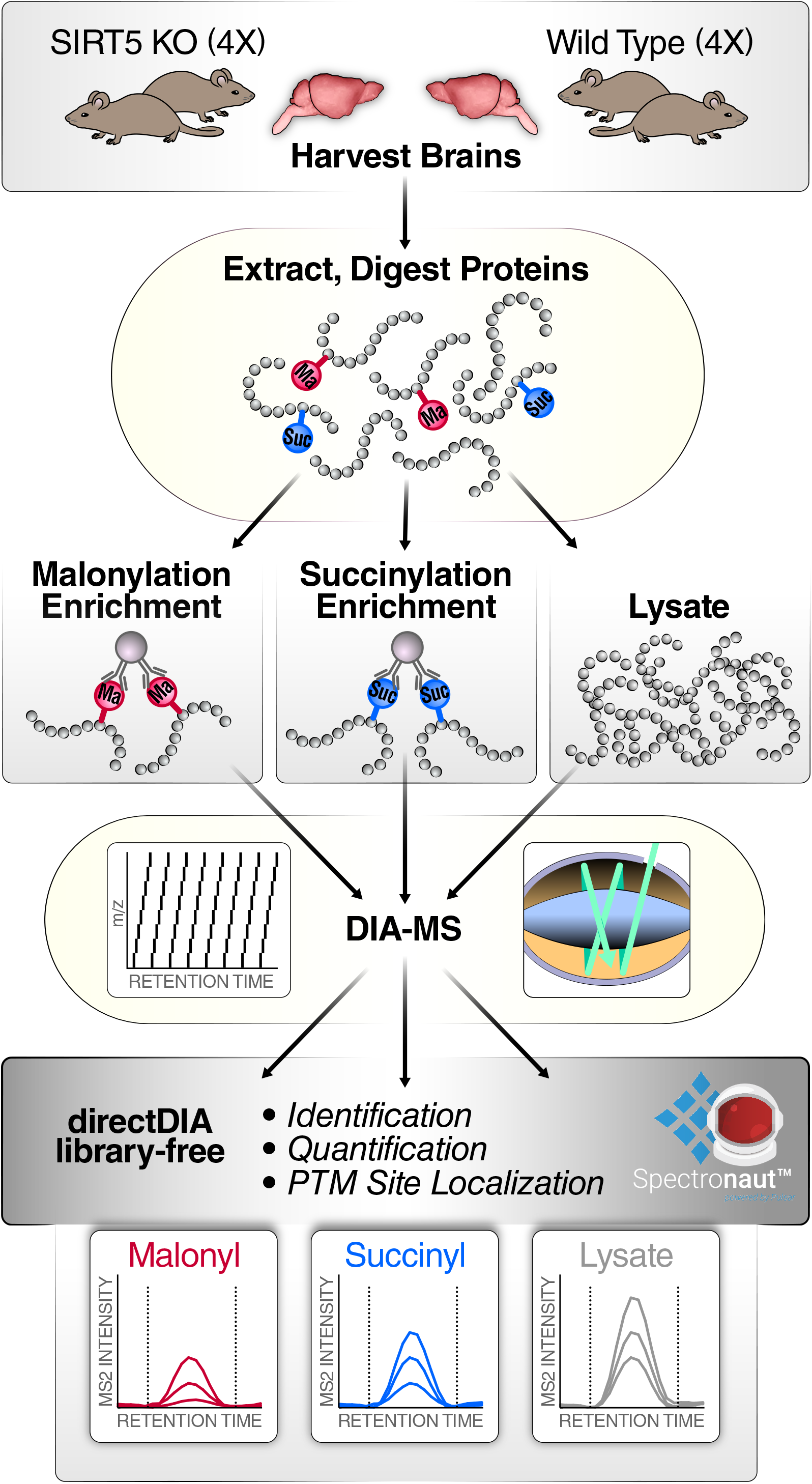
Workflow for investigating SIRT5-induced malonylome and succinylome remodeling in mouse brain. Biological quadruplicates of wild-type and Sirt5^(-/-)^ mouse brains were prepared for in solution trypsin digestion. Samples were enriched for malonyl (Ma) peptides or succinyl (Suc) peptides using antibody-conjugated beads. PTM-enriched samples and samples resulting from whole-lysate digestion were individually analyzed on a nanoLC-Orbitrap Eclipse Tribrid system operating in data-independent acquisition (DIA) mode. Finally, data were processed with directDIA (Biognosys) using a library-free strategy for identification, quantification, and PTM site localization.

To overcome the semi-stochasticity of DDA and the potential dynamic exclusion of PTM peptide isomers, we explored and refined the capabilities of DIA for PTM analysis. Indeed, DIA collects MS/MS maps of all detectable precursor ions contained in the samples to provide comprehensive identification and reproducible and accurate quantification of peptides. DIA acquisitions were performed on an Orbitrap Eclipse Tribrid platform (for detailed protocol see **Appendix 1**). Each cycle was composed of a MS1 scan, followed by 26 MS2 scans collected using a variably windowed isolation scheme [49], where smaller windows are preferred in highly populated *m/z* regions, and larger windows otherwise (*e*.*g*., at higher m/z). This provides reduced MS/MS spectrum complexity, less interferences and increased specificity. The number of data points defining the chromatographic peaks are an important feature in MS2 XIC-based quantification, and an average value of 13 data points per peak was implemented here, which ensured highly accurate quantification.

DIA data were finally processed using a spectral library-free strategy and the directDIA algorithm embedded in Spectronaut, with refined parameters detailed in **Appendix 1**. The directDIA module uses the same DIA files to generate a spectral library and perform a targeted extraction for PTM identifications, site localization, and relative quantification. While DIA data processing commonly relies on an experimentally generated spectral library from prior DDA acquisitions [47], this library-free strategy represents an asset for PTM peptide analysis by reducing the amount of material and acquisition time needed. One DIA acquisition per sample is sufficient to build the spectral library and to perform the relative and accurate quantification. Moreover, using the identical DIA data to generate the spectral library and perform the targeted extraction enabled an efficient retention time calibration (**Figure 2A**). Small retention time windows of a median value less than 5 min from the library retention time were used for the precursor and fragment ion extraction steps, greatly improving and refining the specificity of the assay. This directDIA workflow of mouse WT and SIRT5-KO brains (**Figure 1**) resulted in the identification and quantification of a total of 466 malonylated peptides, corresponding to 423 unique lysine malonylation sites, from 241 malonylated protein groups (**Figure 2B**; **Table S3**), and 2,211 succinylated peptides, corresponding to 1,867 unique lysine succinylation sites, from 561 succinylated protein groups (**Figure 2B**; **Table S4**).

**FIGURE 2.**
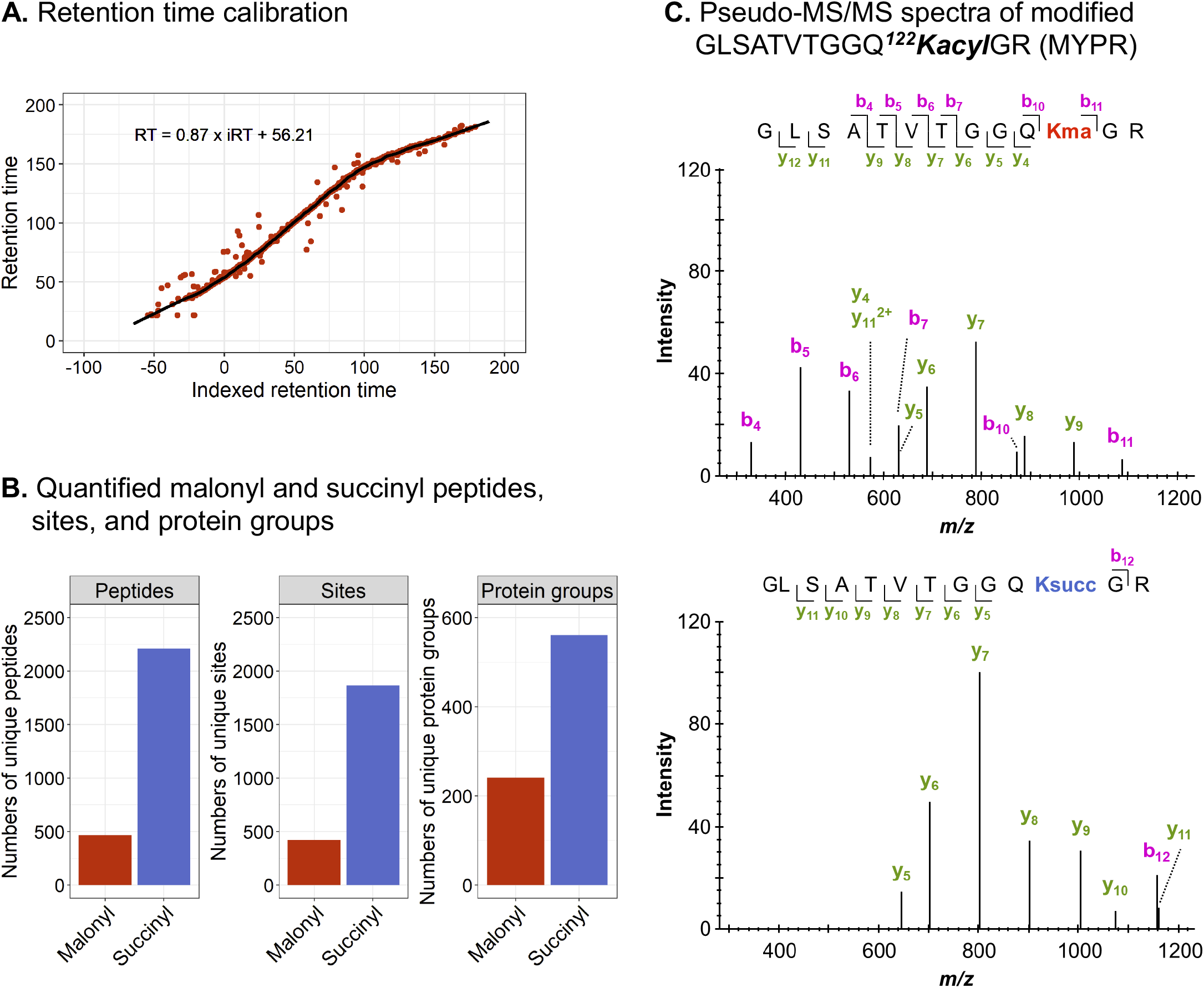
Performance of the workflow for PTM sample analysis. (A) Retention time calibration obtained for a replicate of the malonyl data set. Red dots correspond to peptides used for the calibration, and the black line to the non-linear calibration curve. (B) Number of unique malonylated (red) and succinylated (blue) peptides, sites, and protein groups quantified. (C) Pseudo-MS/MS spectra generated by Spectronaut for the malonylated (top) and succinylated (bottom) peptide GLSATVTGGQ^**122**^**K**GR of the myelin proteolipid protein (MYPR). Detected y ions show similar fragmentation patterns for both malonylated and succinylated peptides.

In addition to PTM identification, PTM site localization is particularly crucial and, at the same time, challenging. To determine the PTM position within the peptide sequence, the data processing software Spectronaut applies a DIA-specific PTM site localization score that assesses site-confirming fragment ions and site-refuting fragment ions for each specific PTM site [61] (also see Biognosys documentation). In addition, the algorithm provides sub-scores that assess the isotopic distribution of fragment ions (note: often singly-charged fragment ions have a limited isotopic envelope), removal of interfering fragment ions, mass accuracy and intensity of the fragment ions [39, 61]. The distribution of the maximum lysine acylation site probability obtained from the four SIRT5-KO replicates and four WT replicates revealed that 84% (390/466) of the malonylated peptides and 95% (2100/2211) of the succinylated peptides achieved a site probability above 0.99 (**Figure S1A-B**), which indicates highly confident PTM site localization.

High confidence in PTM site localization is indeed a cornerstone for assessing PTM crosstalk and differentiating PTM isomers. As an example, **Figure 2C** displays the pseudo-MS/MS spectra obtained for the peptide GLSATVTGGQ^**122**^**K**GR of the myelin proteolipid protein (MYPR), and the lysine residue K122 was identified as both malonylated and succinylated, as evidenced by differentiating ions: y_4_ to y_12_. Moreover, this localization scoring algorithm in Spectronaut allowed us to precisely differentiate 31 pairs of succinyl site isomers and two pairs of malonyl site isomers. For example, two succinylation sites, K242 and K248, were identified from the same peptide sequence, G^**242**^**K**PDVVV^**248**^**K**EDEEEYKR of the mitochondrial acetyl-CoA acetyltransferase (THIL) (**Figure S2**).

To understand the role of malonylation and succinylation in the brain, the subcellular localizations of the quantified malonylated and succinylated proteins were determined. After removing proteins for which no organelle information could be retrieved (six succinylated and two malonylated proteins), 189 malonylated and 433 succinylated proteins were included in the analysis. The proteins were mapped to various subcellular localizations, mainly mitochondrial, cytosolic, plasma membrane and nuclear (**Figure S3A-B**). This is in line with the reported localizations of SIRT5 itself, which is mostly present in the mitochondria, and also in the cytosol and nucleus [23, 24]. In the mouse brains, 71% (308) of the succinylated proteins and 43% (81) of the malonylated proteins were mitochondrial. Yang *et al*. reported a similar proportion of succinylated proteins in human AD brains [62]. However, and more interestingly, the proportions of proteins mapped to other subcellular localizations varied, depending on the PTM type. At least 23% of the malonylated proteins were detected in each cytosolic, plasma membrane and nuclear compartments, against only 19% maximum of the succinylated proteins. IceLogo heatmaps were then generated to determine the preference of each amino acid residue apart from the modified lysine residues, considering sequences 15 amino acids long (**Figure 1C-D**). Alanine, glycine and lysine residues had a high probability of being found upstream and downstream of the modified lysine for the malonylated and succinylated sequences, whereas serine and proline residues had low probabilities of being near the modified lysine.

Overall, these results show that the PTM-DIA workflow combining PTM affinity-based enrichment, comprehensive DIA acquisitions and refined library-free directDIA data processing is highly efficient for investigating and quantifying acylomes from tissues, as here from the brain. DDA-based quantification, such as DDA-TMT, DDA-SILAC and DDA-LFQ, is often used to quantify PTM peptides, but it is biased toward abundant peptides, it presents limited accuracy, and it can potentially lead to PTM isomer exclusion when using dynamic exclusion. Here, DIA allows for the generation of highly comprehensive and accurately quantifiable maps of the PTM samples. More importantly, library-free DIA data analysis is particularly relevant as it is very easy to implement, compatible with low amount of material, reduces the acquisition time, and overcomes issues of DDA PTM analysis. The directDIA workflow assesses all possible PTM site combinations that, combined to the powerful PTM site localization scoring algorithm of Spectronaut, offers highly confident PTM site localization.

### 3.2. Brain malonylome and succinylome remodeling by SIRT5

Acylation groups are specifically removed from lysine residues by lysine deacylases or sirtuins [16]. SIRT5, a member of the NAD^+^-dependent deacylase sirtuin family, has demalonylase [11, 26, 27] and desuccinylase [11, 24] activities (**Figure 3A**). To explore the mouse brain malonylome and succinylome, we compared the changes in PTM peptide levels between WT and SIRT5-KO mice, where SIRT5-KO is expected to result in increased malonylation and succinylation levels. Supervised clustering analysis using partial least squares-discriminant analysis revealed a clear separation between the two groups, explaining 44% and 56% of the variability for the malonyl (**Figure 3C**) and succinyl (**Figure 3D**) datasets, respectively. **Figure 3E-F** displays the volcano plots of the SIRT5-induced PTM changes. For the brain malonylome, 171 peptides, corresponding to 164 unique malonylation sites, were significantly altered with 169 upregulated malonyl peptides and two downregulated malonyl peptides (**Figure 3E-F**; **Supplementary Table S3**), when comparing SIRT5-KO to WT mouse brains. The top five upregulated malonyl peptides showed Log_2_(fold-change) above 16, and belong to the mitochondrial adenylate kinase 4 (KAD4) with site K45, the mitochondrial succinate-semialdehyde dehydrogenase (SSDH) with site K382, band 4.1-like protein 2 (E41L2) with site K188, stromal membrane-associated protein 1 (SMAP1) with site K12, and gephyrin (GEPH) with site K324. In the brain succinylome, 640 succinylated peptides, corresponding to 578 unique succinylation sites, were significantly altered, including 630 upregulated succinyl peptides and 10 downregulated succinyl peptides (**Figure 3F**; **Table S4**), when comparing SIRT5-KO to WT mouse brains. The top five upregulated succinylated sites respectively belong to the following proteins: mitochondrial enoyl-CoA delta isomerase 2 (ECI2) with site K60, mitochondrial acetyl-CoA acetyltransferase (THIL) with sites K260 and K263, mitochondrial ATP synthase-coupling factor 6 (ATP5J) with site K79, mitochondrial ATP synthase subunit O (ATPO) with site K73, and acyl-coenzyme A thioesterase 13 (ACO13) with site K17. It is worth noting that SIRT5 induced large remodeling of the brain malonylome and succinylome, but also remarkably strong fold-changes, with Log_2_(fold-change) up to 21, and median Log_2_(fold-change) of 3.7 and 8.0, respectively. This reflects and emphasizes the fact that the corresponding modified peptides were barely or not at all detected in the WT mouse brains.

**FIGURE 3.**
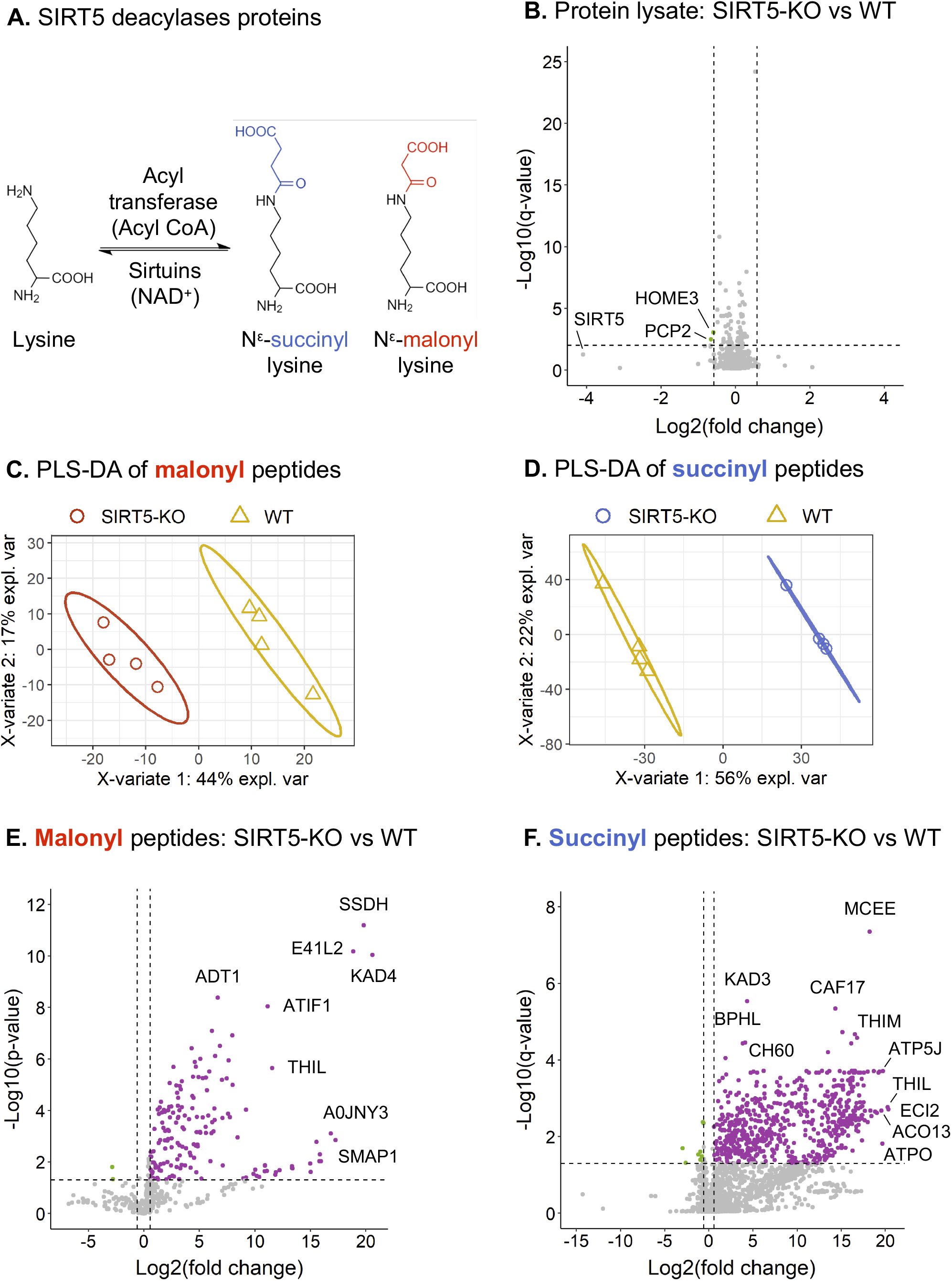
Remodeling of mouse brain malonylome and succinylome by SIRT5. (A) Schematic of the regulation of lysine malonylation and succinylation. (B) Volcano plot of the 2,837 protein groups quantified in whole-lysate samples, with two downregulated proteins groups obtained for SIRT5-KO vs WT comparison. (C-D) Supervised clustering analysis using partial least squares-discriminant analysis (PLS-DA) performed on all malonylated (C) and succinylated (D) peptides quantified in the wild-type (WT; yellow) and SIRT5-KO (red for malonyl, blue for succinyl) samples. (E) Volcano plot of the 466 quantified malonylated peptides, with 169 up- and two downregulated malonylated peptides obtained for SIRT5-KO vs WT comparison. (F) Volcano plot of the 2,211 quantified succinylated peptides, with 630 up- and 10 downregulated succinylated peptides obtained for SIRT5-KO vs WT comparison.

Similar to the subcellular localization observed for all quantified malonylated and succinylated proteins, SIRT5-regulated proteins were mostly mitochondrial in both cases: 49 (52%) for malonylated proteins, and 186 (92%) for the succinylated proteins, with different proportions among the compartments between the PTM types, similarly to was observed for all quantified modified proteins (**Figure S3C-D**).

Additionally, the protein lysate analysis revealed that out of the 2,837 protein groups quantified with at least two unique peptides, only two protein groups were significantly altered (**Figure 3B**; **Table S5**), in SIRT5-KO vs WT mouse brains. The two changing proteins, homer protein homolog 3 (HOME3) and Purkinje cell protein 2 (PCP2), were significantly downregulated. As expected, SIRT5 showed the strongest downregulation fold-change, close to a significant q-value threshold. Altogether, these protein level results demonstrate that SIRT5 very marginally affects the proteome, but instead leads to a large remodeling of the malonylome and succinylome of mouse brains.

### 3.3. Crosstalk between brain lysine malonylation and succinylation

To further explore SIRT5-induced brain malonylome and succinylome remodeling, we investigated the cross-talk between these two PTMs, as they both occur on lysine residues. **Figure 4A** displays the distribution of the SIRT5-targeted lysine sites. For each modification, most proteins had single modification sites, representing 72% (89) of the SIRT5-targeted proteins for malonylation, and 51% (126) for succinylation. More precisely, in this study, up to four malonyl sites per protein were identified, with the isoform 4 of myelin basic protein (P04370-4) presenting four malonyl groups at K57, K98, K166 and K180. The myelin basic protein presents a large number of isoforms, resulting from alternative splicing and PTMs, among which the “classic” isoforms 4-13 are major components of the oligodendrocyte myelin membrane in the central nervous system. They interact with various proteins, including actin, tubulin, calmodulin and several SH3-domain proteins [63, 64]. The multiple malonylation of the isoform 4 may impact its interaction with negatively charged proteins, altering its roles in myelin events. Regarding succinylation, 89% (222) of the SIRT5-targeted proteins showed up to 4 modified sites, 2.0% (5) presenting more than 10 modified sites, with a maximum number of 15 different sites as obtained for the mitochondrial trifunctional enzyme subunit alpha (ECHA). The mitochondrial trifunctional enzyme is a protein complex involved in mitochondrial fatty acid beta-oxidation pathway, however its hypersuccinylation has been reported not to affect its activity [65].

**FIGURE 4.**
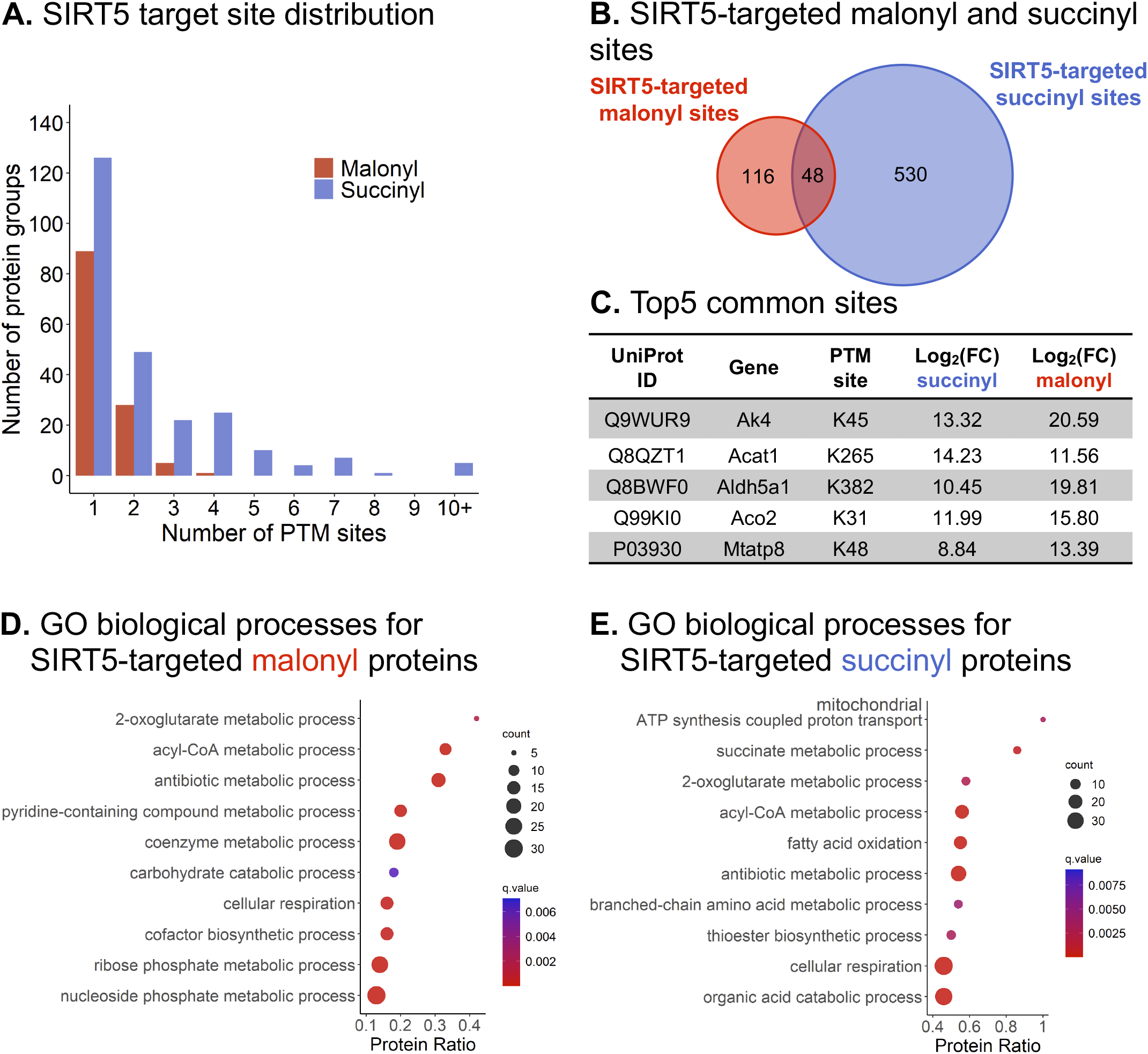
Crosstalk between brain lysine malonylation and succinylation. (A) Distribution of the malonyl (red) and succinyl (blue) sites targeted by SIRT5. (B) Venn diagram comparing the SIRT5-targeted malonyl and succinyl sites. The 48 common sites belong to 36 protein groups, and the top 5 sites are listed (C). (D-E) Dotplots showing the ConsensusPathDB [53, 54] gene ontology (GO) biological processes enriched for SIRT5-regulated malonyl (D) and succinyl (E) proteins.

Interestingly, 48 SIRT5-targeted sites, corresponding to 36 succinylated/malonylated protein groups, were identified both malonylated and succinylated in our dataset (**Figure 4B**; **Supplementary Table S6**). For 30 sites, the succinyl SIRT5-KO vs WT ratios were higher than the malonyl ratios. These proteins, which are mostly mitochondrial, are involved in various metabolic processes, such as ketone body metabolic process, succinate metabolic process, acyl-CoA metabolic process, 2-oxoglutarate metabolic process, and cellular respiration (**Table S7**). As an example, the top 5 common sites are K45 of the mitochondrial adenylate kinase 4 (KAD4), K265 of the mitochondrial acetyl-CoA acetyltransferase (THIL), K382 of the mitochondrial succinate-semialdehyde dehydrogenase (SSDH), K31 of the mitochondrial aconitate hydratase (ACON), and K48 of the ATP synthase protein 8 (ATP8) (**Figure 4C**). The two non-mitochondrial proteins are the myelin proteolipid protein (MYPR; site K122, which is part of a cytoplasmic domain), and the isoform 4 of myelin basic protein (MBP; site K98), two major components of myelin.

### 3.4. Acylation occurs mostly on proteins related to metabolism and neurodegeneration

To determine the biological processes and pathways enriched with SIRT5-KO, over-representation analyses were performed using ConsensusPathDB database [53, 54]. GO analysis showed that various metabolism-related processes were upregulated for both malonyl and succinyl datasets (**Figure 4D-E**; **Table S7**). More particularly, the 2-oxoglutarate metabolic process, acyl-CoA metabolic process, and cellular respiration were commonly enriched, whereas mitochondrial ATP synthesis coupled proton transport, succinate metabolic process, and fatty acid oxidation were specific to the succinyl data. KEGG pathway enrichment analyses revealed that numerous metabolism-related pathways, including amino acid metabolism, carbohydrate metabolism, lipid metabolism and oxidative phosphorylation, were commonly enriched (**Figure 5A-B**; **Table S7**). Nervous system-related pathways, such as GABAergic synapse and synaptic vesicle cycle for malonyl dataset, and retrograde endocannabinoid signaling for succinyl data set, were also enriched. More interestingly, SIRT5 succinylome remodeling affected several neurodegenerative disease-related pathways, including Alzheimer’s disease (AD), amyotrophic lateral sclerosis (ALS), Huntington’s disease (HD), prion disease, and Parkinson’s disease (PD). The latter was also affected by SIRT5 malonylome remodeling.

**FIGURE 5.**
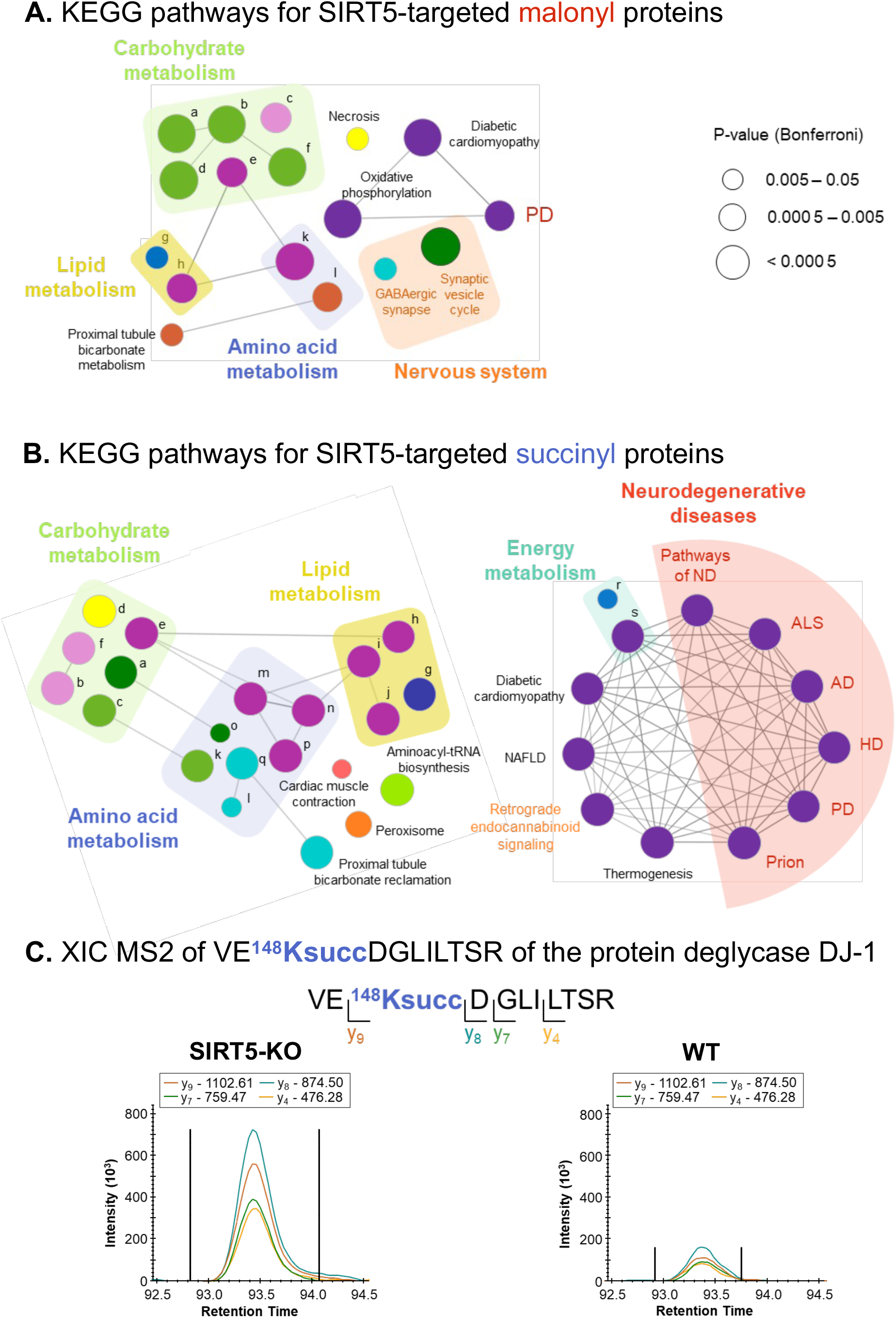
SIRT5 affects proteins involved in Parkinson’s disease. (A-B) ClueGO [56] KEGG pathway enrichment and network analysis of SIRT5-regulated malonyl (A) and succinyl (B) proteins. Pathway-connecting edges represent Kappa score above 40%. Same color pathways have at least 50% similarity. Pathways are listed in **Table S7E**. Neurogenerative diseases: Pathways of ND, pathways of neurodegeneration; ALS, amyotrophic lateral sclerosis; AD, Alzheimer’s disease; HD, Huntington’s disease; PD, Parkinson’s disease; Prion, prion disease. NAFLD, non-alcoholic fatty liver disease. (C) Extracted ion chromatograms of VE^**148**^**Ksucc**DGLILTSR (m/z 665.86, z = 2+) of the protein deglycase DJ-1 (PARK7).

### 3.5. Parkinson’s disease pathway is enriched in SIRT5-regulated malonylome and succinylome

The PD pathway is enriched for both SIRT5-regulated malonylome and succinylome, and so, we further investigated proteins associated with this pathway. This concerned 10 malonylated proteins, and 33 succinylated proteins (**Table S8**), among which eight proteins were common: the ADP/ATP translocases 1 and 2 (ADT1/ANT1 and ADT2/ANT2, respectively), the NADH dehydrogenase [ubiquinone] 1 alpha subcomplex subunit 7 (NDUA7), the mitochondrial cytochrome b-c1 complex subunit 2 (QCR2), the subunits alpha, beta and O of the mitochondrial ATP synthase (ATPA, ATPB and ATPO, respectively), and the mitochondrial ATP synthase F(0) complex subunit B1 (AT5F1). More particularly, four SIRT5-targeted PTM sites were commonly malonylated and succinylated: K498 of ATPA, K233 of AT5F1, K40 of NDUA7, and K147 of ADT1/ANT1 (**Figure 5**; **Figure S4**).

Parkinson’s disease is the second most common and a complex multifactorial neurological disorder which is characterized by parkinsonism, the accumulation of intracytoplasmic Lewy bodies and Lewy neurites – mostly corresponding to α-synuclein aggregates, and loss of dopaminergic neurons in the substantia nigra pars compacta [66]. Mutations of several genes, including SNCA, LRRK2, PINK1, PARKIN and DJ-1, are causes of the different forms of PD.

Mitochondrial dysfunction plays a crucial role in neurodegeneration in PD [66, 67]. We observed that all five complexes of the respiratory chain were highly acylated and SIRT5 targeted Complexes 1 and 5 primarily. Indeed, a total of 34 sites from 13 subunits of the NADH dehydrogenase or Complex 1, involved in transferring electrons from NADH to the respiratory chain, were succinylated. Among these, one site from NDUA7 showed both malonylation and succinylation (**Figure S4**). Mitochondrial ATP synthase or Complex V, responsible for the production of ATP in mitochondria, was highly acylated. Among the components reported associated with PD from the KEGG enrichment analysis, eight subunits were malonylated and/or succinylated, with 22 total sites targeted by SIRT5: ATPA (4 sites), ATPB (4 sites), ATPG (1 site) and ATP5E (2 sites) of the F_1_ domain, and ATP5H (1 site), ATP5J (4 sites), ATPO (5 sites) and AT5F1 (1 site) of the F_0_ domain. More particularly, the site K498 of the subunit alpha (ATPA) and the site K233 of the F(0) complex subunit B1 (AT5F1) showed dual malonylation and succinylation (**Figure S4**). Recently, DJ-1 was reported to bind ATPB [68] and to regulate its synthesis and position to allow proper inner mitochondrial membrane permeability, hence acting as an important player in the metabolism of dopaminergic neurons.

The protein DJ-1, encoded by the Park7 gene, protects dopamine neurons from cell death in PD [69, 70]. DJ-1 acts as an oxidative stress sensor and protects mitochondria against oxidative stress, an important mediator in PD. Moreover, it presents a chaperone function: it prevents oligomerization of α-synuclein. DJ-1 is also involved in protective responses that depend on transcription factors, including p53. DJ-1 mutations cause PD autosomal recessive forms [66], leading to its loss of function, however, through an unclear mechanism. Caspase-6 (CASP6) cleaves DJ-1 at D149, and the D149A mutation impairs CASP6 proteolysis, abolishing its protective role associated with p53-dependent cell death [71]. Interestingly, K148 adjacent to the CASP6 cleavage site was highly succinylated in SIRT5-KO mouse brain (**Figure 5C**). Further investigations would need to be conducted to investigate whether this modification impairs CASP6 cleavage.

The ADP/ATP translocase 1 (ADT1/ANT1), encoded by the Slc25a4 gene, is the principal exchanger of cytosolic ADP and mitochondrial ATP across the inner mitochondrial membrane, providing this metabolite to the mitochondrial ATP synthase, and is involved in the maintenance of various mitochondrial functions, including parkin-mediated mitophagy [72]. More particularly, the mechanism involves the interaction of ANT1 with the TIM23 complex, the presequence translocase of the inner membrane, that mediates the import of proteins containing a cleavage presequence into the mitochondrial matrix, and more specifically with TIM44, a matrix-sided component of TIM23. ANT1 mutation G146E/K147D prevented its interaction with TIM44, impairing mitophagy regulation. This mutation changes the net charge of this two-residue sequence from positive to negative, as both glutamic acid and aspartic acid residues are negatively charged. Interestingly, lysine malonylation or succinylation, that we observed at site K147 (**Figure S4**), changes the positive charge of the lysine residue to negative, which also potentially impacts the interaction of ANT1 with TIM44. ANT1 interacts with several other proteins to form the mitochondrial permeability transition pore (mPTP). In PD, lower levels of ANT1 leads to mPTP being open, triggering a cascade of cell apoptosis program. Moreover, ANT1 co-aggregates with α-synuclein, potentially causing neuronal damages [73].

The calcium/calmodulin-dependent protein kinase type II (Camk2) is a Ca^2+^/calmodulin-dependent synapse-enriched kinase that is involved in the regulation of numerous biological and neuronal processes, including Ca^2+^ regulation, synaptic plasticity and gene expression. Camk2 is linked to PD via Ca^2+^ signaling dysregulation in dopamine neurons [74]. Here, the subunit beta of the calcium/calmodulin-dependent protein kinase type II (KCC2B) is reported malonylated on K33 (**Figure S4**), between its close ATP binding region in position 20-28 and ATP binding site in position 43.

Lastly, the protein β-synuclein (SYUB) was not reported as part of the KEGG mouse Parkinson’s disease pathway. However, it has a neuroprotective function as its interaction with α-synuclein inhibits α-synuclein aggregation, which limits PD progression [75, 76]. β-synuclein is composed of three domains, the N-terminal domain, the central non-amyloid-component (NAC) domain and the C-terminal domain, and all three of them are required to inhibit α-synuclein fibrilization. Further investigations are needed to assess the impact of the malonylation of the sites K23 and K34, located in the N-terminal region of β-synuclein (**Figure S4**) on its interaction with other synucleins.

## 4. Concluding Remarks

Here, we presented an efficient and straightforward workflow for PTM identification, site localization and quantification using PTM affinity-based enrichment, comprehensive DIA acquisitions and spectral library-free DIA data processing with refined parameters. We have demonstrated the high performances of directDIA software for generating a spectral library, retrieving identification as well as robust and highly accurate quantification information from the same DIA data in a fully automated manner. Moreover, the PTM localization site algorithm embedded into Spectronaut enables confident PTM site localization and thus precise isomer differentiation. Plus, in directDIA, all possible PTM sites combinations are considered and the identification of PTM sites is not limited to the modified peptides contained in the spectral library, enabling more comprehensive profiling of PTM peptides.

This strategy offers the possibility to comprehensively, systematically and quantitatively profile hundreds modified peptides, while limiting the amount of starting material as well as the acquisition time. While applied to acylation modifications, we believe that this workflow can be applied to any modification types of interest.

Finally, our study provides deep insights into the remodeling of mouse brain malonylome and succinylome upon SIRT5 regulation. Indeed, 171 malonylated sites from 123 protein groups and 640 succinylated sites from 249 protein groups were targeted by SIRT5, while protein lysate was barely affected. These acylated proteins localized mainly in mitochondria, although malonylated proteins were also significantly present in the cytosol, nucleus and at the plasma membrane. We revealed 46 common SIRT5-regulated malonylation and succinylation sites from 36 protein groups, suggesting potential PTM crosstalk. Carbohydrate, lipid amino acid and energy metabolisms, and nervous system-related pathways were affected by SIRT5. Moreover, several pathways related to neurodegenerative diseases were enriched, which is supported by the known roles of SIRT5 in these diseases [30], including PD. More particularly, SIRT5 targeted the five complexes of the respiratory chain and the protein DJ-1/Park7 – related to oxidative damage, a hallmark of PD, ANT1/Slc25a4 – involved in mitophagy, the subunit beta of Camk2 – important for Ca^2+^ regulation in dopaminergic neurons, and β-synuclein – showing a neuroprotective action.

Liu *et al*. [77] reported that SIRT5 protects the degeneration of dopaminergic neurons induced by the neurotoxin 1-methyl-4-phenyl-1,2,3,6-tetrahydropyridine (MPTP) in the substantia nigra pars compacta by preserving the expression level of superoxide dismutase 2 (SOD2). This antioxidant enzyme is involved in scavenging reactive oxygen species (ROS) in mitochondria and thus prevents oxidative stress. Similarly, in our study, SIRT5 did not regulate expression of the dismutase SOD2; however, we identified five acylation sites: K68, K114 and K108 were succinylated, and K122 and K108 were dually malonylated and succinylated. MPTP inhibits the mitochondrial Complex 1, which alters energy production and leads to ROS production. Interestingly, we showed that 34 acylation sites from 13 subunits of Complex 1 – all succinylated and one with both succinylation and malonylation – were regulated by SIRT5.

Baeken *et al*. [78] reported that oxidative stress induced SIRTs autophagic degradation in dopaminergic neurons in *in vitro* PD models. The authors hypothesized that the hyper-acetylation of the proteome upon SIRT depletion could have a protective role against oxidative damage by favoring acetylation-mediated transcriptional activation and preventing lysine cross-linking with other amino acid residues or nucleic acids.

Understanding the mechanisms underlying protein lysine malonylation and succinylation in metabolic diseases or neurodegenerative diseases is highly relevant and present a high biomedical interest for the discovery of novel therapeutic targets against these diseases. We are providing an efficient DIA-PTM workflow that uses directDIA to accurately quantify acylation level changes in a sample-preserving approach, that also favors high throughput analysis.

## Supporting information

Appendix_Protocol

## Abbreviations

AD: Alzheimer’s disease
ADP: Adenosine diphosphate
ALS: Amyotrophic lateral sclerosis
ATP: Adenosine triphosphate
CoA: Coenzyme A
DDA: data-dependent acquisition
DIA: data-independent acquisition
FA: formic acid
HD: Huntington’s disease
IAP: Immunoaffinity purification
KEGG: Kyoto Encyclopedia of Genes and Genomes
KO: knock-out
mPTP: Mitochondrial permeability transition pore
NAD: Nicotinamide adenine dinucleotide
PD: Parkinson’s disease
PLS-DA: Partial least squares-discriminant analysis
SIRT5: Sirtuin 5
TEAB: Triethylammonium bicarbonate
WT: wild-type
XIC: extracted ion chromatogram

## 5. Associated Data

Raw data and complete MS data sets have been uploaded to the Center for Computational Mass Spectrometry, to the MassIVE repository at UCSD, and can be downloaded using the following link: https://massive.ucsd.edu/ProteoSAFe/dataset.jsp?task=0d9cfcca844340988e6e772d8444dd99 (MassIVE ID number: MSV000089606; ProteomeXchange ID: PXD034275).

[Note to the reviewers: To access the data repository MassIVE (UCSD) for MS data, please use: Username: MSV000089606_reviewer; Password: winter].

## Acknowledgments

We acknowledge the support of instrumentation for the Orbitrap Eclipse Tribrid system from the NCRR shared instrumentation grant 1S10 OD028654 (PI: Birgit Schilling).

