## Supplementary Figure S1 for "In-depth Analysis of the Sirtuin 5-regulated Mouse Brain Acylome using Library-free Data-Independent Acquisitions"

### Slide 1
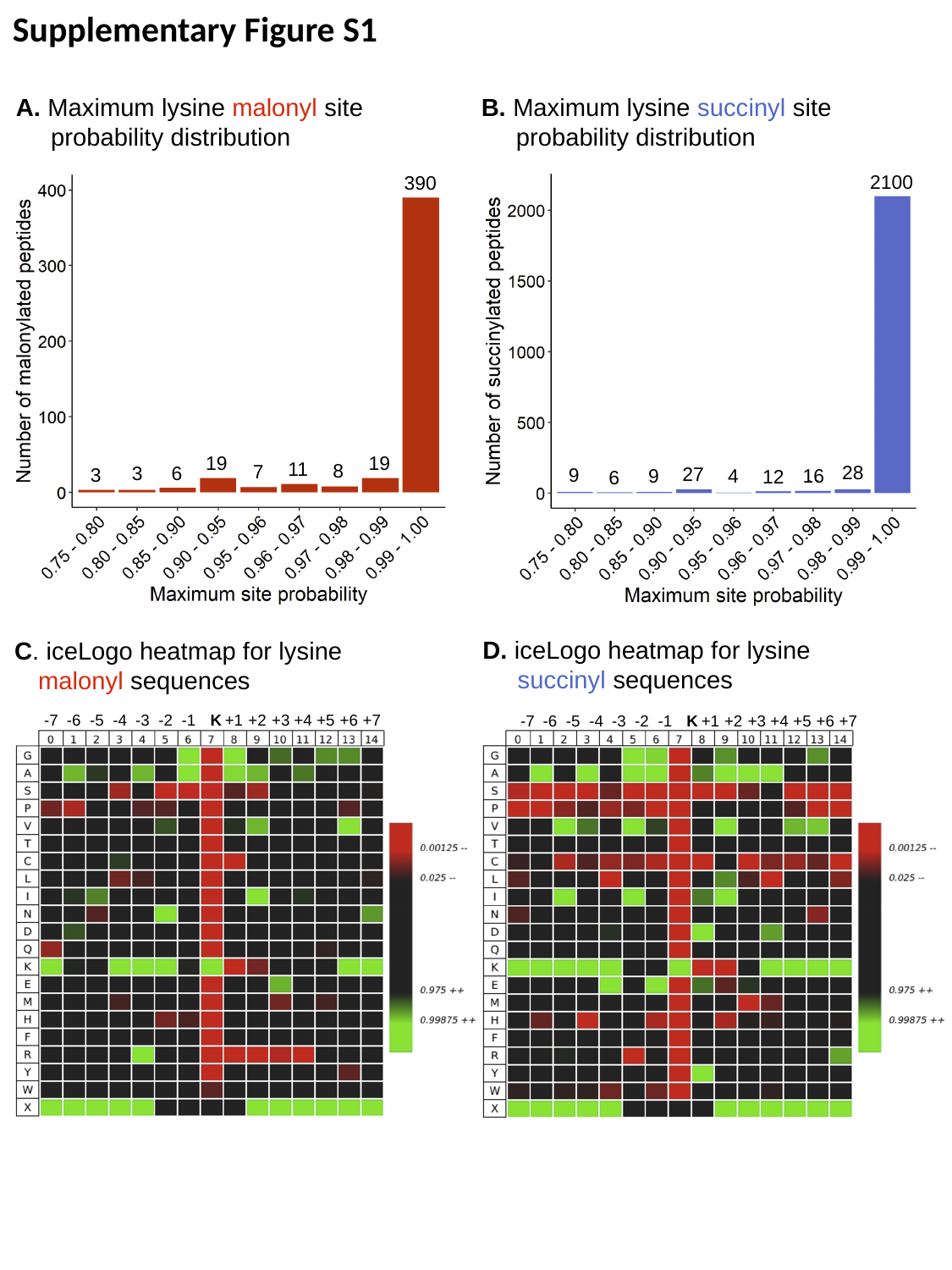

Supplementary Figure S1
B. Maximum lysine succinyl site
 probability distribution
A. Maximum lysine malonyl site
 probability distribution
2100
28
27
9
16
9
4
12
6
390
19
19
11
8
7
6
3
3
D. iceLogo heatmap for lysine
 succinyl sequences
C. iceLogo heatmap for lysine malonyl sequences
-7
-6
-5
-4
-3
-2
-1
K
+1
+2
+3
+4
+5
+6
+7
-7
-6
-5
-4
-3
-2
-1
K
+1
+2
+3
+4
+5
+6
+7

### Slide 2
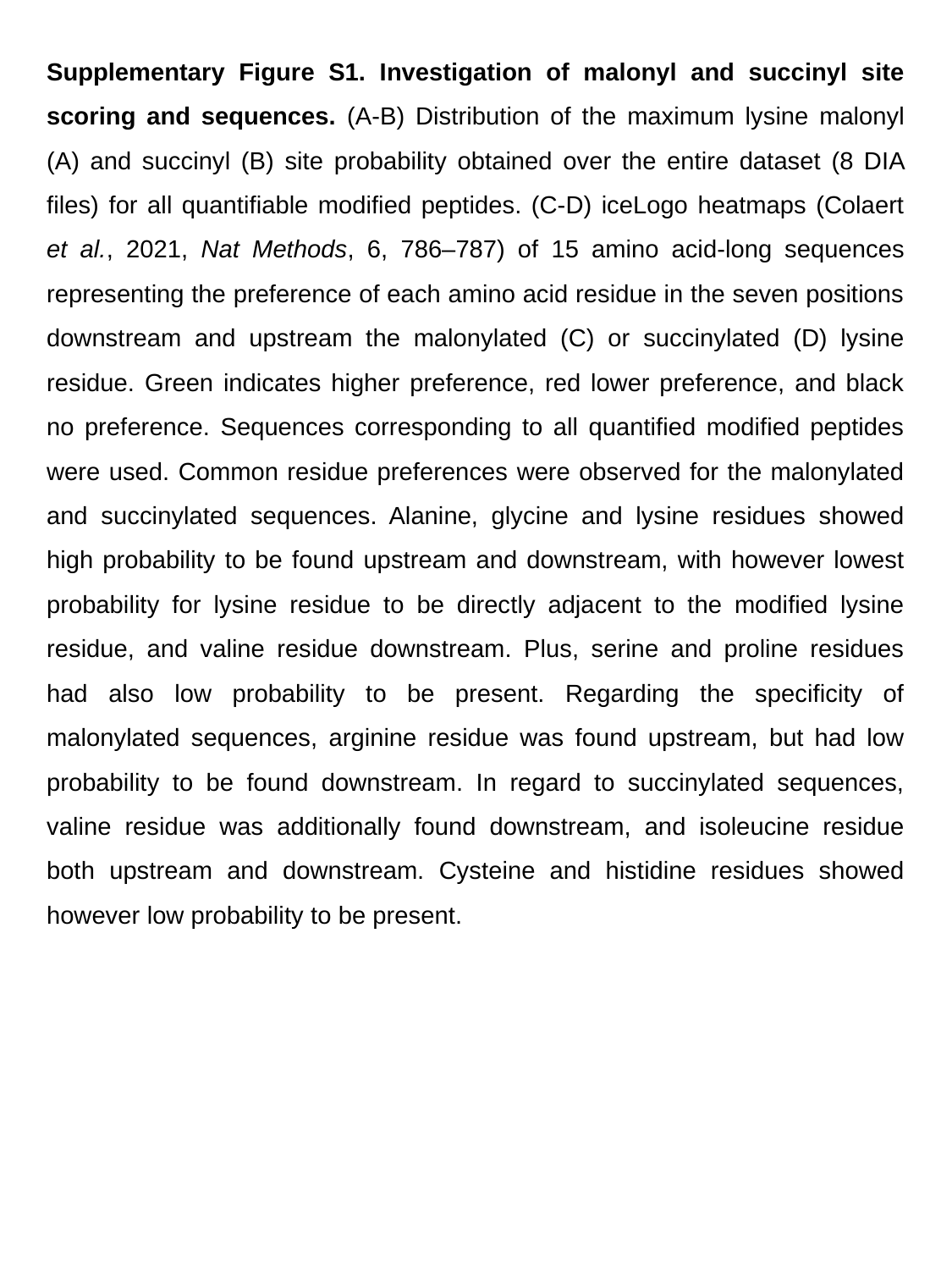

Supplementary Figure S1. Investigation of malonyl and succinyl site scoring and sequences. (A-B) Distribution of the maximum lysine malonyl (A) and succinyl (B) site probability obtained over the entire dataset (8 DIA files) for all quantifiable modified peptides. (C-D) iceLogo heatmaps (Colaert et al., 2021, Nat Methods, 6, 786–787) of 15 amino acid-long sequences representing the preference of each amino acid residue in the seven positions downstream and upstream the malonylated (C) or succinylated (D) lysine residue. Green indicates higher preference, red lower preference, and black no preference. Sequences corresponding to all quantified modified peptides were used. Common residue preferences were observed for the malonylated and succinylated sequences. Alanine, glycine and lysine residues showed high probability to be found upstream and downstream, with however lowest probability for lysine residue to be directly adjacent to the modified lysine residue, and valine residue downstream. Plus, serine and proline residues had also low probability to be present. Regarding the specificity of malonylated sequences, arginine residue was found upstream, but had low probability to be found downstream. In regard to succinylated sequences, valine residue was additionally found downstream, and isoleucine residue both upstream and downstream. Cysteine and histidine residues showed however low probability to be present.
