## Supplementary Figure S2 for "In-depth Analysis of the Sirtuin 5-regulated Mouse Brain Acylome using Library-free Data-Independent Acquisitions"

### Slide 1
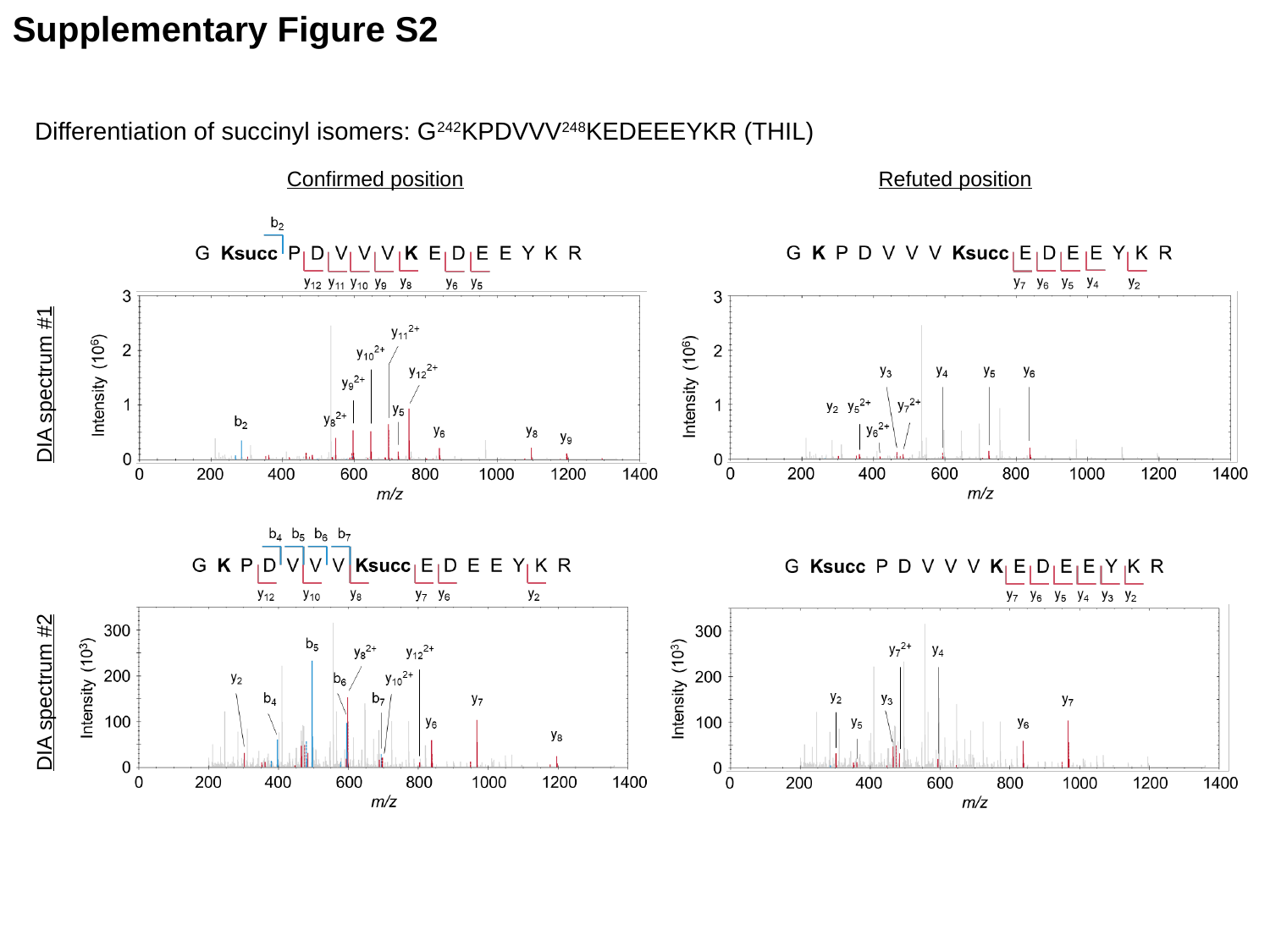

Supplementary Figure S2
Differentiation of succinyl isomers: G^242^KPDVVV^248^KEDEEEYKR (THIL)
Refuted position
Confirmed position
DIA spectrum #1
DIA spectrum #2

### Slide 2
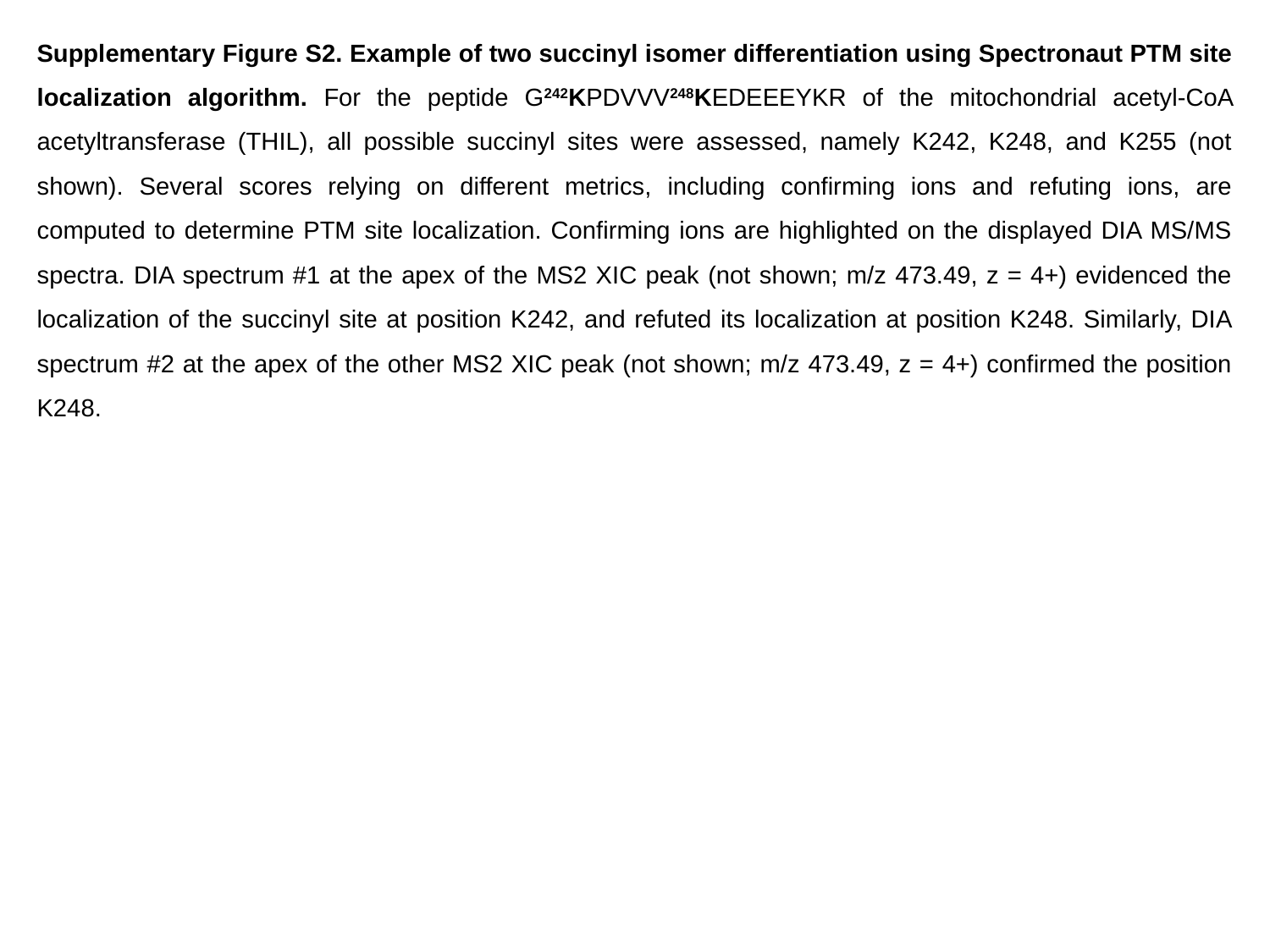

Supplementary Figure S2. Example of two succinyl isomer differentiation using Spectronaut PTM site localization algorithm. For the peptide G^242^KPDVVV^248^KEDEEEYKR of the mitochondrial acetyl-CoA acetyltransferase (THIL), all possible succinyl sites were assessed, namely K242, K248, and K255 (not shown). Several scores relying on different metrics, including confirming ions and refuting ions, are computed to determine PTM site localization. Confirming ions are highlighted on the displayed DIA MS/MS spectra. DIA spectrum #1 at the apex of the MS2 XIC peak (not shown; m/z 473.49, z = 4+) evidenced the localization of the succinyl site at position K242, and refuted its localization at position K248. Similarly, DIA spectrum #2 at the apex of the other MS2 XIC peak (not shown; m/z 473.49, z = 4+) confirmed the position K248.
