## Supplementary Figure S3 for "In-depth Analysis of the Sirtuin 5-regulated Mouse Brain Acylome using Library-free Data-Independent Acquisitions"

### Slide 1
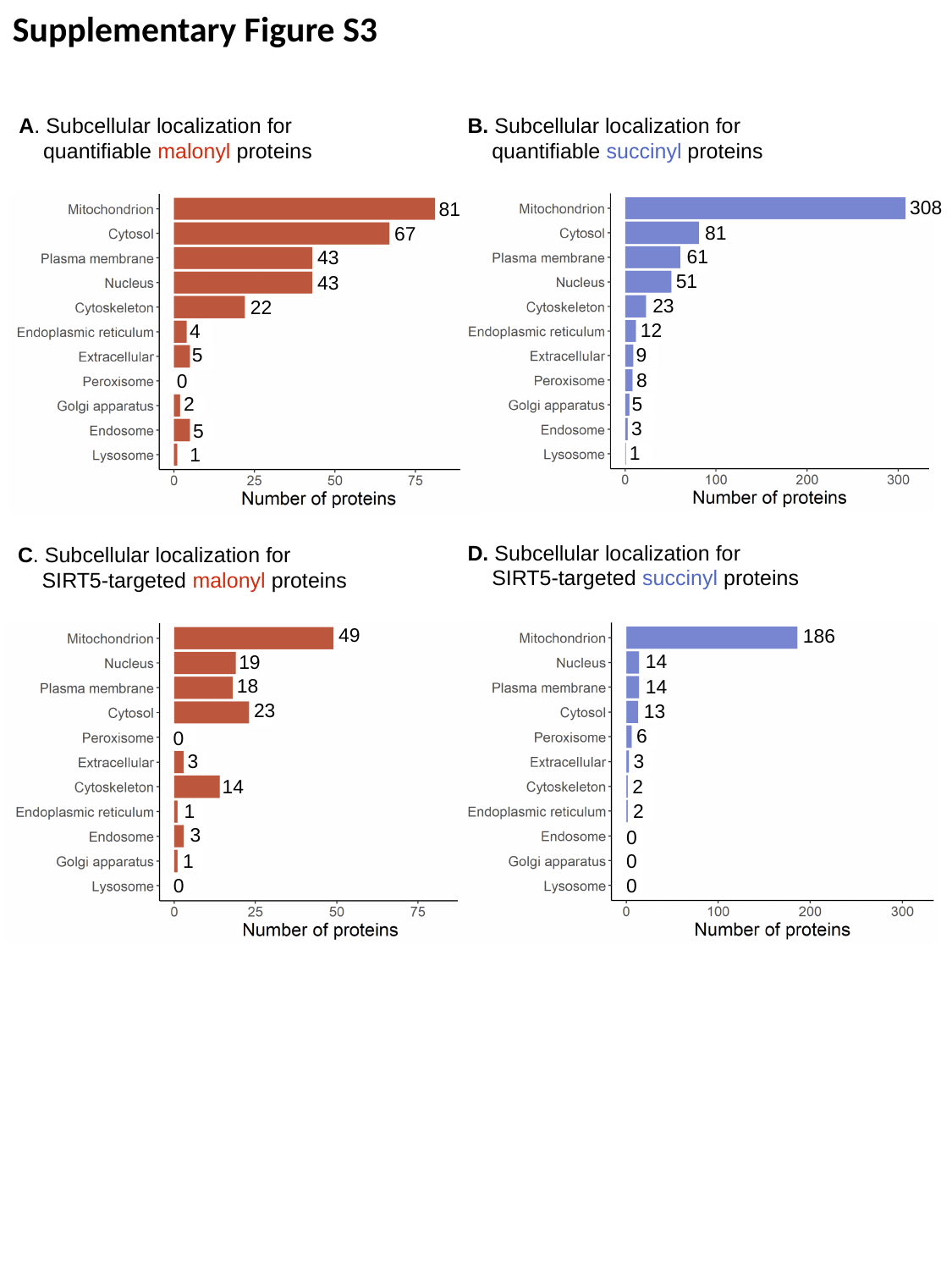

Supplementary Figure S3
A. Subcellular localization for quantifiable malonyl proteins
B. Subcellular localization for quantifiable succinyl proteins
308
81
61
51
23
12
9
8
5
3
1
81
67
43
43
22
4
5
0
2
5
1
D. Subcellular localization for
SIRT5-targeted succinyl proteins
C. Subcellular localization for
SIRT5-targeted malonyl proteins
49
19
18
23
0
3
14
1
3
1
0
186
14
14
13
6
3
2
2
0
0
0

### Slide 2
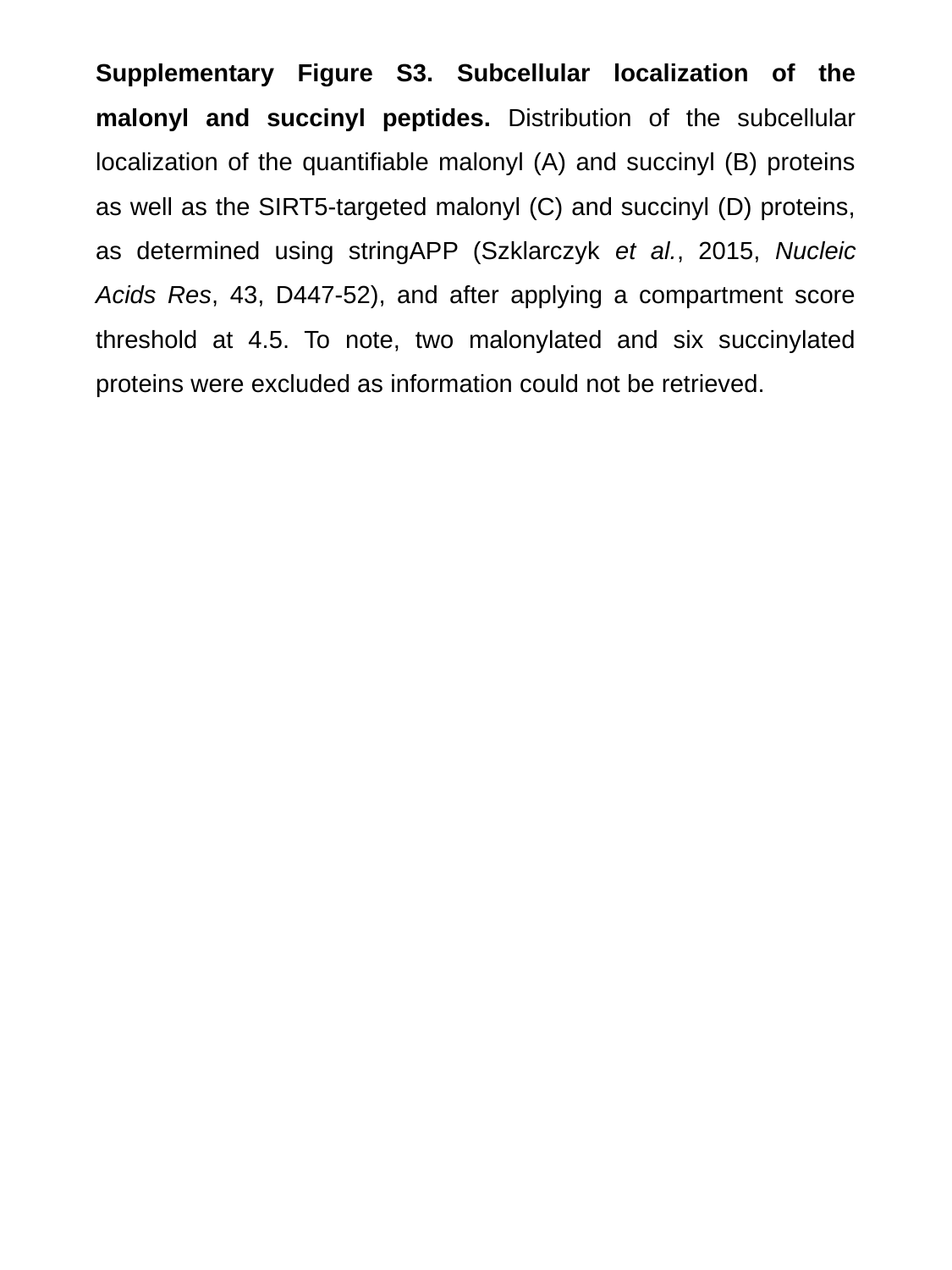

Supplementary Figure S3. Subcellular localization of the malonyl and succinyl peptides. Distribution of the subcellular localization of the quantifiable malonyl (A) and succinyl (B) proteins as well as the SIRT5-targeted malonyl (C) and succinyl (D) proteins, as determined using stringAPP (Szklarczyk et al., 2015, Nucleic Acids Res, 43, D447-52), and after applying a compartment score threshold at 4.5. To note, two malonylated and six succinylated proteins were excluded as information could not be retrieved.
