## Supplementary Figure S4 for "In-depth Analysis of the Sirtuin 5-regulated Mouse Brain Acylome using Library-free Data-Independent Acquisitions"

### Slide 1
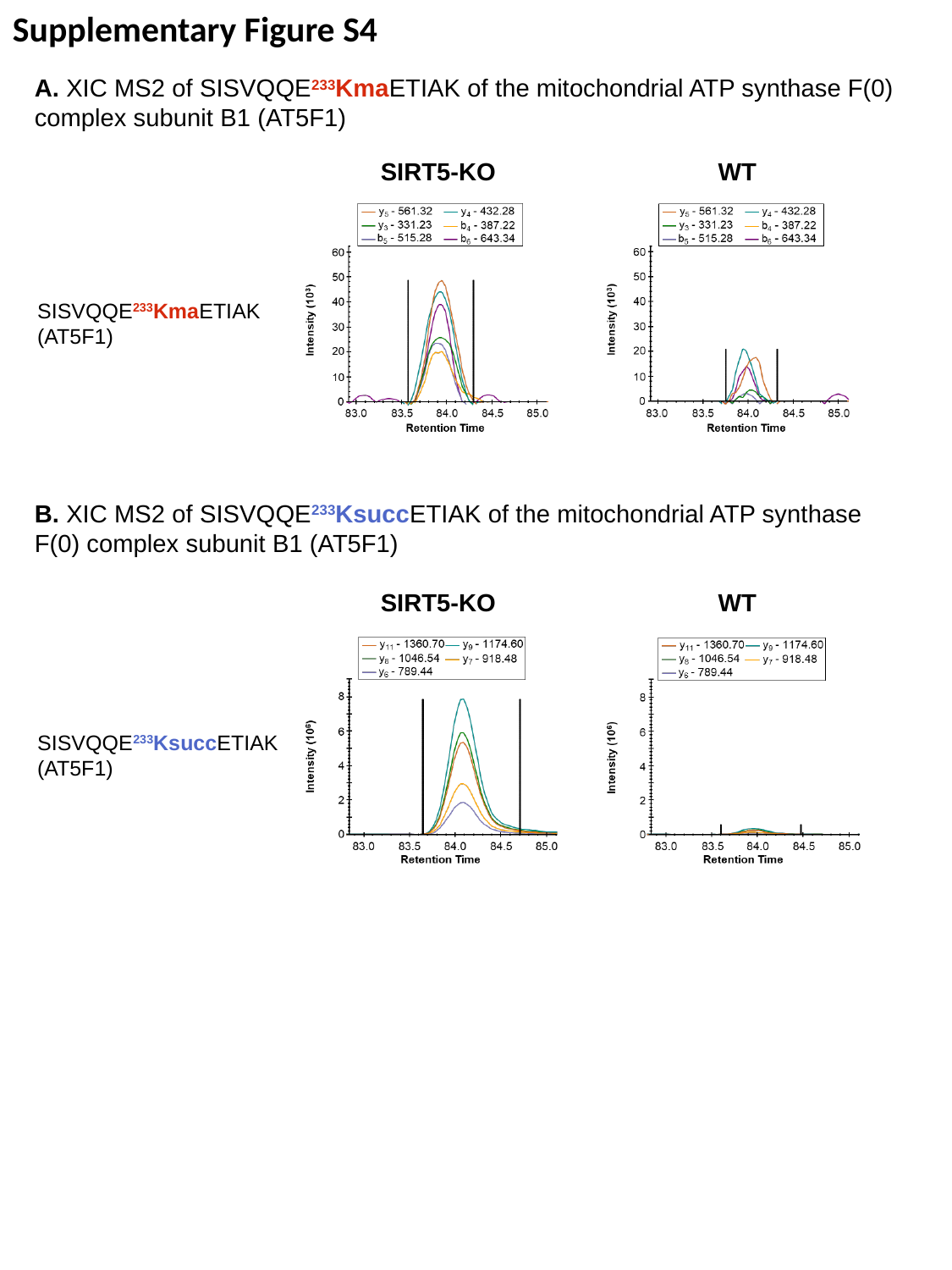

Supplementary Figure S4
A. XIC MS2 of SISVQQE^233^KmaETIAK of the mitochondrial ATP synthase F(0) complex subunit B1 (AT5F1)
SIRT5-KO
WT
SISVQQE^233^KmaETIAK
(AT5F1)
B. XIC MS2 of SISVQQE^233^KsuccETIAK of the mitochondrial ATP synthase F(0) complex subunit B1 (AT5F1)
SIRT5-KO
WT
SISVQQE^233^KsuccETIAK
(AT5F1)

### Slide 2
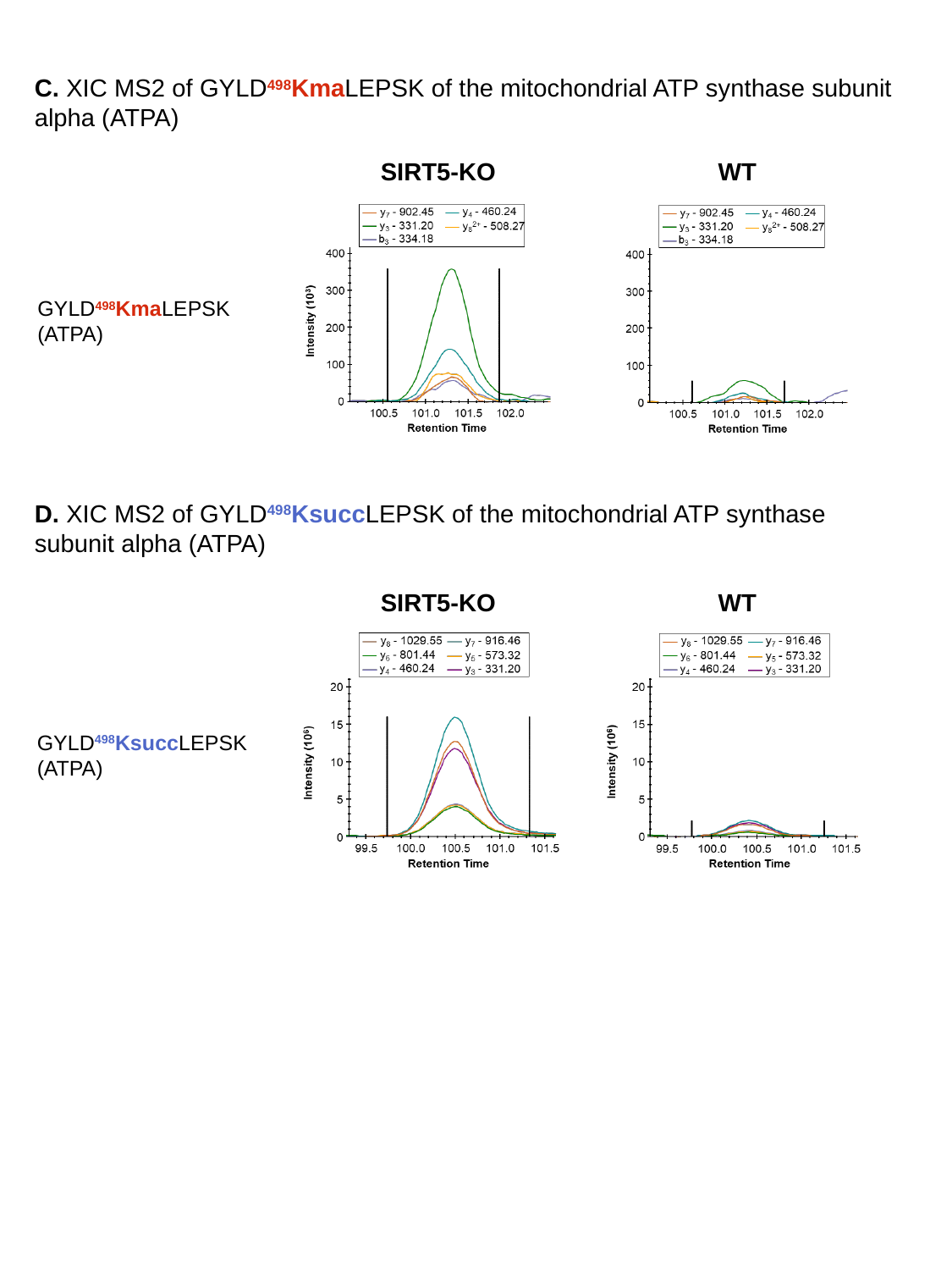

C. XIC MS2 of GYLD^498^KmaLEPSK of the mitochondrial ATP synthase subunit alpha (ATPA)
SIRT5-KO
WT
GYLD^498^KmaLEPSK
(ATPA)
D. XIC MS2 of GYLD^498^KsuccLEPSK of the mitochondrial ATP synthase subunit alpha (ATPA)
SIRT5-KO
WT
GYLD^498^KsuccLEPSK
(ATPA)

### Slide 3
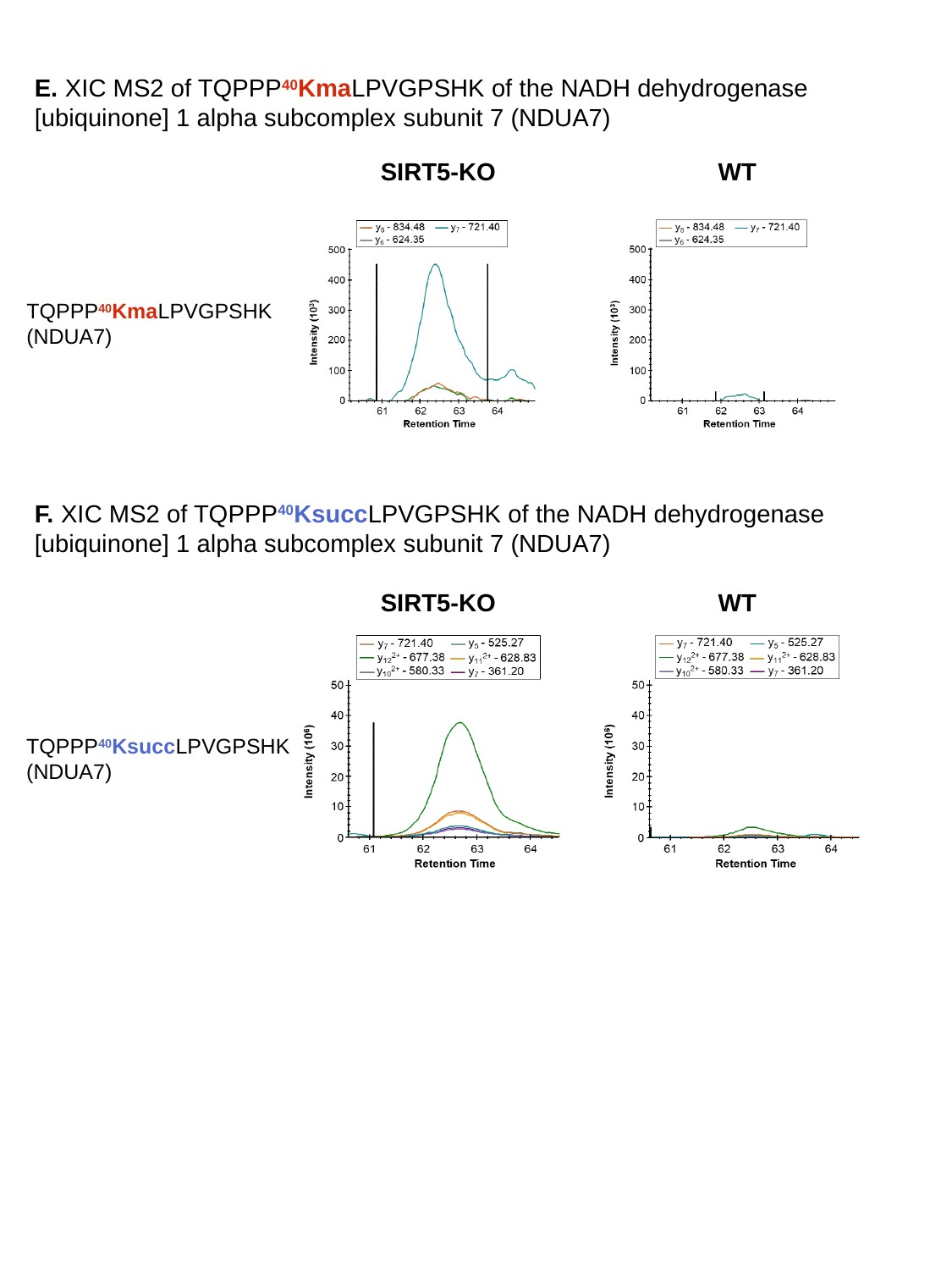

E. XIC MS2 of TQPPP^40^KmaLPVGPSHK of the NADH dehydrogenase [ubiquinone] 1 alpha subcomplex subunit 7 (NDUA7)
SIRT5-KO
WT
TQPPP^40^KmaLPVGPSHK
(NDUA7)
F. XIC MS2 of TQPPP^40^KsuccLPVGPSHK of the NADH dehydrogenase [ubiquinone] 1 alpha subcomplex subunit 7 (NDUA7)
SIRT5-KO
WT
TQPPP^40^KsuccLPVGPSHK
(NDUA7)

### Slide 4
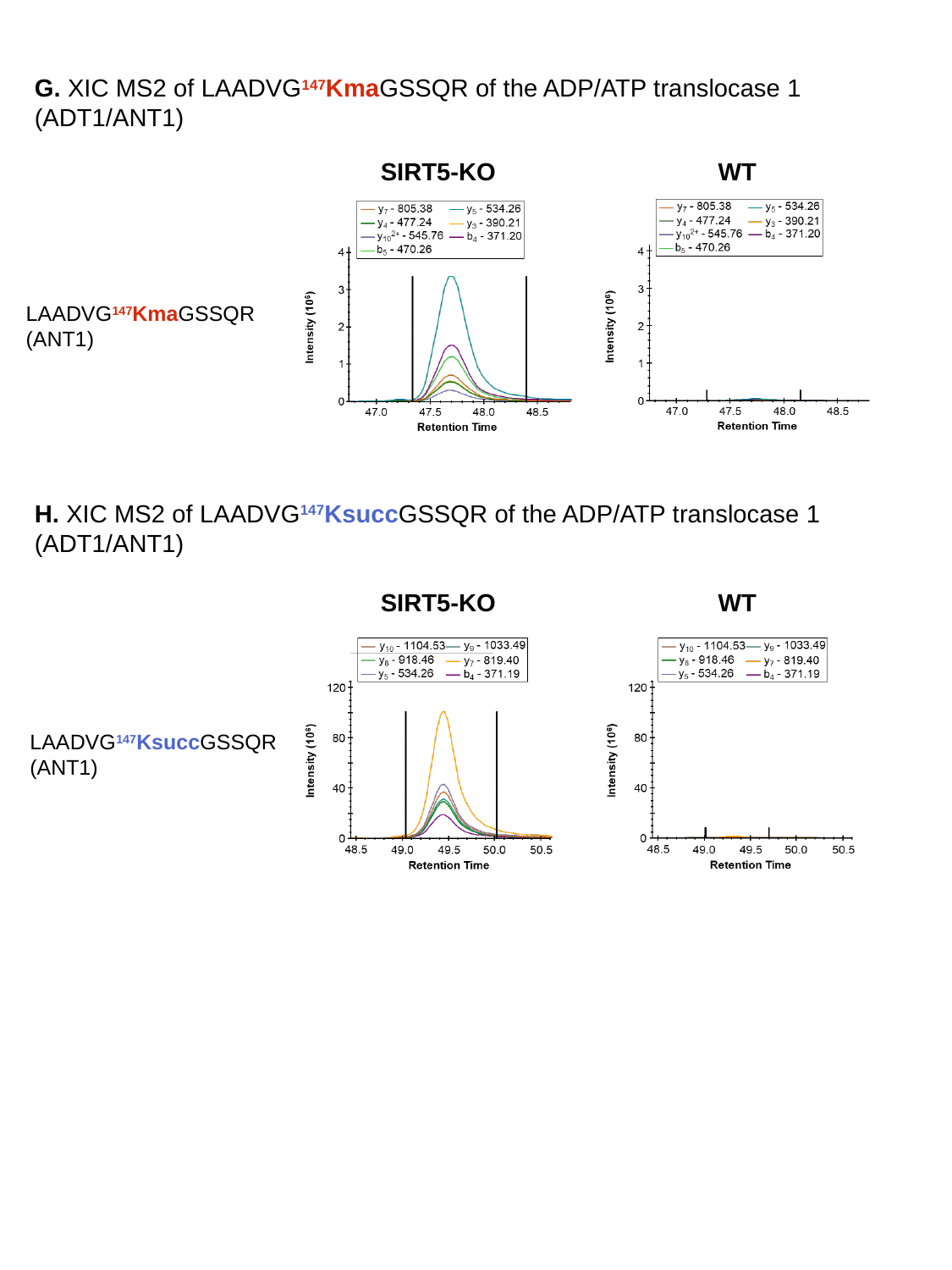

G. XIC MS2 of LAADVG^147^KmaGSSQR of the ADP/ATP translocase 1 (ADT1/ANT1)
SIRT5-KO
WT
LAADVG^147^KmaGSSQR
(ANT1)
H. XIC MS2 of LAADVG^147^KsuccGSSQR of the ADP/ATP translocase 1 (ADT1/ANT1)
SIRT5-KO
WT
LAADVG^147^KsuccGSSQR
(ANT1)

### Slide 5
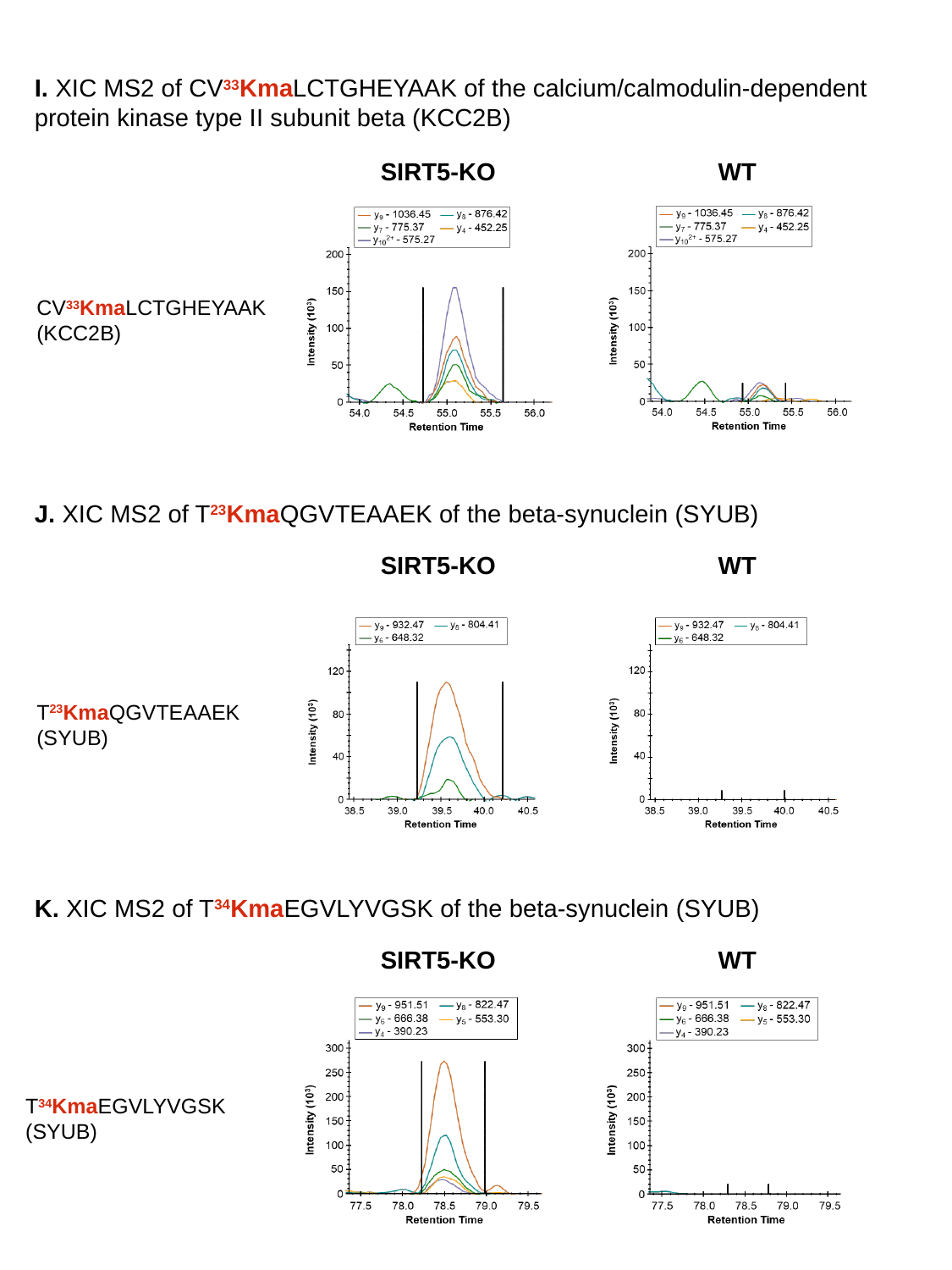

I. XIC MS2 of CV^33^KmaLCTGHEYAAK of the calcium/calmodulin-dependent protein kinase type II subunit beta (KCC2B)
SIRT5-KO
WT
CV^33^KmaLCTGHEYAAK
(KCC2B)
J. XIC MS2 of T^23^KmaQGVTEAAEK of the beta-synuclein (SYUB)
SIRT5-KO
WT
T^23^KmaQGVTEAAEK
(SYUB)
K. XIC MS2 of T^34^KmaEGVLYVGSK of the beta-synuclein (SYUB)
SIRT5-KO
WT
T^34^KmaEGVLYVGSK
(SYUB)

### Slide 6
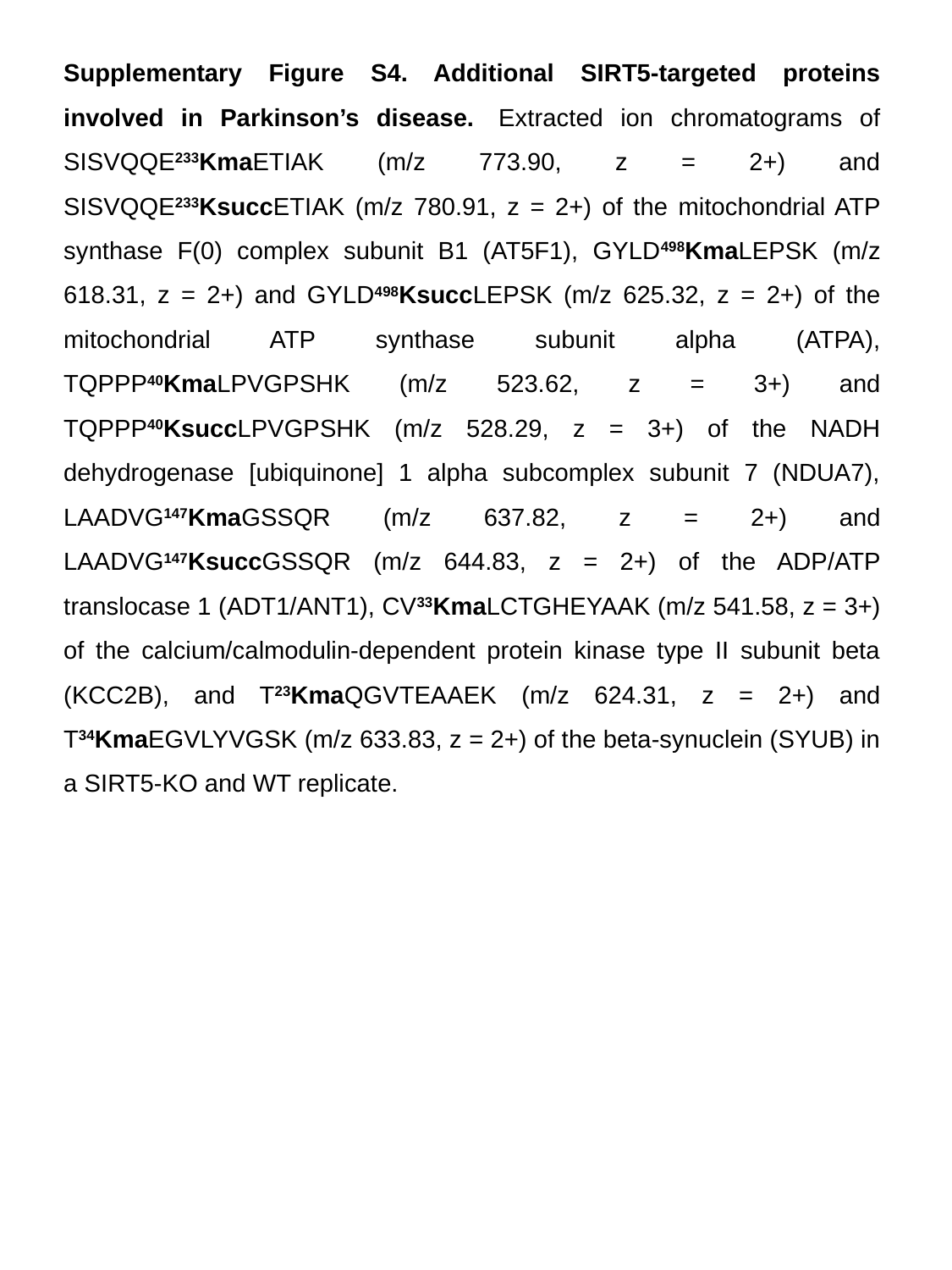

Supplementary Figure S4. Additional SIRT5-targeted proteins involved in Parkinson’s disease.  Extracted ion chromatograms of SISVQQE^233^KmaETIAK (m/z 773.90, z = 2+) and SISVQQE^233^KsuccETIAK (m/z 780.91, z = 2+) of the mitochondrial ATP synthase F(0) complex subunit B1 (AT5F1), GYLD^498^KmaLEPSK (m/z 618.31, z = 2+) and GYLD^498^KsuccLEPSK (m/z 625.32, z = 2+) of the mitochondrial ATP synthase subunit alpha (ATPA), TQPPP^40^KmaLPVGPSHK (m/z 523.62, z = 3+) and TQPPP^40^KsuccLPVGPSHK (m/z 528.29, z = 3+) of the NADH dehydrogenase [ubiquinone] 1 alpha subcomplex subunit 7 (NDUA7), LAADVG^147^KmaGSSQR (m/z 637.82, z = 2+) and LAADVG^147^KsuccGSSQR (m/z 644.83, z = 2+) of the ADP/ATP translocase 1 (ADT1/ANT1), CV^33^KmaLCTGHEYAAK (m/z 541.58, z = 3+) of the calcium/calmodulin-dependent protein kinase type II subunit beta (KCC2B), and T^23^KmaQGVTEAAEK (m/z 624.31, z = 2+) and T^34^KmaEGVLYVGSK (m/z 633.83, z = 2+) of the beta-synuclein (SYUB) in a SIRT5-KO and WT replicate.
