## Supplementary Information/Methods for "In-depth Analysis of the Sirtuin 5-regulated Mouse Brain Acylome using Library-free Data-Independent Acquisitions"

Corresponding author:

**Materials and Methods**

**Chemicals**

LC-MS-grade acetonitrile and water were obtained from Burdick & Jackson (Muskegon, MI). Reagents for protein chemistry, including urea, iodoacetamide (IAA), dithiothreitol (DTT), triethylammonium bicarbonate (TEAB), and formic acid (FA) were purchased from Sigma-Aldrich (St. Louis, MO). Sequencing-grade trypsin was purchased from Promega (Madison, WI). HLB Oasis SPE cartridges were purchased from Waters (Milford, MA).

**Mouse Brains**

Animal study was performed according to protocols approved by IACUC (the Institutional Animal Care and Use Committee). SIRT5 knock-out (KO) mice were obtained from the Jackson Laboratory (Strain#012757). Mice were housed (12-h light/dark cycle, 22°C) and given unrestricted access to water. Brain tissues were collected from 18 months old females, after 6 hours of refeeding following 24 hours of fasting.

**Protein Digestion and Desalting**

Mouse brain tissues were collected from 2 different conditions with 4 biological replicates each: i) wild-type (WT, n=4), ii) SIRT5^(-/-)^ (SIRT5-KO, n=4). Frozen brains were homogenized in lysis buffer containing 8 M urea, 50 mM Tris, pH 7.5, 7.5, 1x HALT protease inhibitor cocktail (Thermo Fisher Scientific, Waltham, MA), 150 mM NaCl, 5 μM trichostatin A, and 5 mM nicotinamide, and homogenized for 2 cycles with a Bead Beater TissueLyser II (Qiagen, Germantown, MD) at 25 Hz for 3 min each. Lysates were clarified by spinning at 16,500 x *g* for 15 min at 4°C, and the supernatant containing the soluble proteins was collected. Protein concentrations were determined using a Bicinchoninic Acid Protein (BCA) Assay (Thermo Fisher Scientific, Waltham, MA), and subsequently 3-mg and 5-mg protein from each sample were aliquoted for malonyl and succinyl peptide analysis, respectively. Proteins were reduced with 20 mM DTT for 30 min at 37 °C, and after cooling to room temperature, alkylated with 40 mM IAA for 30 min in the dark at room temperature. Samples were diluted 4-fold with 50 mM TEAB, pH 7.5, and proteins were digested overnight with a solution of sequencing-grade trypsin in 50 mM TEAB at a 1:50 (wt:wt) enzyme:protein ratio at 37°C. This reaction was quenched with 1% FA and the sample was clarified by centrifugation at 2,000 x *g* for 10 min at room temperature. Clarified peptide samples were desalted with Oasis 10-mg Sorbent Cartridges. 100 μg of each peptide elution were aliquoted for analysis of protein-level changes, after which all desalted samples were vacuum dried. The 100 µg whole lysate aliquots were re-suspended in 0.2% FA in water at a final concentration of 1 µg/µL and stored for MS analysis. The remaining 4.9 mg digests (for malonyl enrichment) and 2.9 mg digests (for succinyl enrichment) were re-suspended in 1.4 mL of immunoaffinity purification (IAP) buffer (Cell Signaling Technology, Danvers, MA) containing 50 mM 4-morpholinepropanesulfonic acid (MOPS)/sodium hydroxide, pH 7.2, 10 mM disodium phosphate, and 50 mM sodium chloride for PTM enrichment. Peptides were enriched for malonylation with anti-malonyl antibody conjugated to agarose beads from the Malonyl-Lysine Motif Kit (Kit #93872), and for succinylation with anti-succinyl antibody conjugated to agarose beads from the Succinyl-Lysine Motif Kit (Kit #13764; both from Cell Signaling Technology, Danvers, MA). This process was performed according to the manufacturer protocol; however, each sample was incubated in half the recommended volume of washed beads. Peptides were eluted from the antibody-bead conjugates with 0.1% trifluoroacetic acid in water and were desalted using C18 stagetips made in-house. Samples were vacuum dried and re-suspended in 0.2% FA in water. Finally, indexed retention time standard peptides (iRT; Biognosys, Schlieren, Switzerland) [1] were spiked in the samples according to manufacturer’s instructions.

**Mass Spectrometric Analysis**

LC-MS/MS analyses were performed on a Dionex UltiMate 3000 system coupled to an Orbitrap Eclipse Tribrid mass spectrometer (both from Thermo Fisher Scientific, San Jose, CA). The solvent system consisted of 2% ACN, 0.1% FA in H_2_O (solvent A) and 98% ACN, 0.1% FA in H_2_O (solvent B). Proteolytic peptides were loaded onto an Acclaim PepMap 100 C18 trap column (0.1 x 20 mm, 5 µm particle size; Thermo Fisher Scientific) for 5 min at 5 µL/min with 100% solvent A. For the protein lysates, an amount of 200 ng was loaded (protein level analysis), and for the enriched malonylated and succinylated peptides (PTM level analysis), 4 µL of each PTM-enriched sample were injected. Peptides were eluted on an Acclaim PepMap 100 C18 analytical column (75 µm x 50 cm, 3 µm particle size; Thermo Fisher Scientific) at 300 nL/min using the following gradient of solvent B: 2% for 5 min, linear from 2% to 20% in 125 min, linear from 20% to 32% in 40 min, and up to 80% in 1 min, with a total gradient length of 210 min.

All samples – protein level analysis and PTM level analyses – were acquired in data-independent acquisition (DIA) mode. Full MS spectra were collected at 120,000 resolution (AGC target: 3e6 ions, maximum injection time: 60 ms, 350-1,650 m/z), and MS2 spectra at 30,000 resolution (AGC target: 3e6 ions, maximum injection time: Auto, NCE: 27, fixed first mass 200 m/z). The isolation scheme consisted in 26 variable windows covering the 350-1,650 m/z range with an overlap of 1 m/z (**Table S1**) [2]. A detailed set-by-step procedure for building the DIA method and subsequent data processing can be found in **Appendix 1**.

Statistical analysis was performed in Skyline, and malonylated peptides with p < 0.05 and absolute fold-change > 1.5 were considered as significantly altered.

**Clustering Analysis**

Partial least squares-discriminant analysis (PLS-DA) of the proteomics data was performed using the package mixOmics [5] in R (version 4.0.2; RStudio, version 1.3.1093).

**Enrichment Analysis**

An over-representation analysis (ORA) was performed using Consensus Path DB-mouse (Release MM11, 14.10.2021) [6, 7] to evaluate which gene ontology (GO) terms were significantly enriched. Gene ontology terms identified from the ORA were subjected to the following filters: q-value < 0.01, and term level > 3 for GO. Dot plots were generated using the ggplot2 package [8] in R (version 4.0.5; RStudio, version 1.4.1106). Kyoto Encyclopedia of Genes and Genomes (KEGG) pathway enrichment analysis was performed using the ClueGO package [9] (version 2.5.8), in Cytoscape [10] (version 3.8.2). Default settings were applied, except that two-sided hypergeometric test Bonferroni-adjusted p-value threshold was set to 0.01. Kappa score threshold was let at 0.4 for drawing pathway-connecting edges. Pathways with the same color indicate at least 50% similarity in genes/term.

**iceLogo Heatmap**

Heatmap of 15 amino acid-long sequences centered around the modified lysine residues were generated using the iceLogo tool [11] to determine the frequency of every amino acid residue around the modified lysine residue. The *Mus musculus* precompiled Swiss-Prot composition was chosen, and the start position was set to 0. The p-value was set at 0.05, and significantly up- and down-regulated residues are colored in shades of green and red, respectively.

**Subcellular Localization**

Subcellular localization was determined using Cytoscape [10] (version 3.8.2) and stringAPP [12] (version 1.7.0), by applying default settings, except that species was defined as *Mus musculus*. Only compartments with scores above 4.5 were considered.
