## Appendix_Protocol for "In-depth Analysis of the Sirtuin 5-regulated Mouse Brain Acylome using Library-free Data-Independent Acquisitions"

**Appendix 1: Bons, J., Rose, J., Zhang, R.; Burton, J. B., Carrico, C., Verdin, E., Schilling, B. In-depth Analysis of the Sirtuin 5-regulated Mouse Brain Acylome using Library-free Data-Independent Acquisitions, *Proteomics***

### Generation of a LC-MS/MS Variable Window Data-independent Acquisition Method for a Thermo Orbitrap Eclipse Platform, and Instructions for Data Processing with directDIA (Spectronaut)

Note: use hyperlinks to go to the individual chapters by clicking on the page numbers above.

#### Chapter 1. Generate a LC-MS/MS method file for variable window DIA

1. Open **XCalibur** and navigate to the **XApps Page**.

    *Note: Here we use XCalibur version 4.3.73.11.*

2. Click **Instrument Setup** to start a new method.

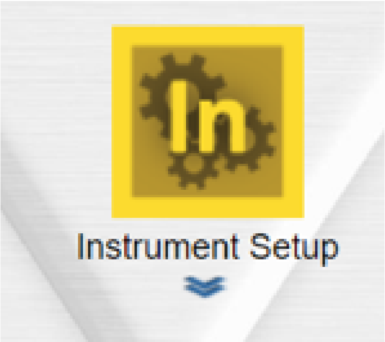

3. Click **Orbitrap Eclipse** to modify the MS method.

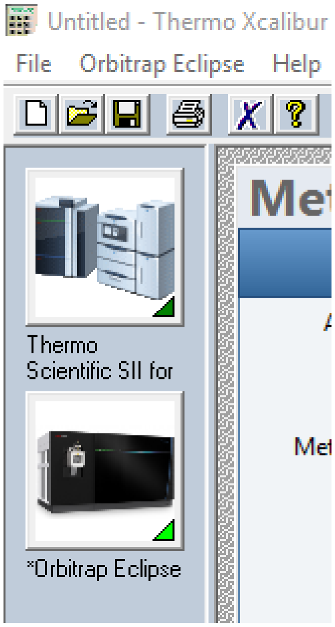

4. Set the **Method Duration** to the length of the chromatographic gradient.

    *Note: In this study, the length of the chromatographic gradient is 210 min.*

5. On the **Settings** menu, change the following parameters:

- set the **Expected LC Peak Width** to 30 s
- check **Advanced Peak Determination**
- set the **Default Charge State** to 3
- set the **Internal Mass Calibration** to Off.

6. On the **Ion Source Properties** menu, change the source conditions to conditions that were optimized for the system, which should be in the following ranges:

- set the **Positive Ion (V)** to 1900 – 2300 V
- set the **Sweep Gas (Arb)** to 0
- set the **Ion Transfer Tube Temp (°C)** to 290 – 310.

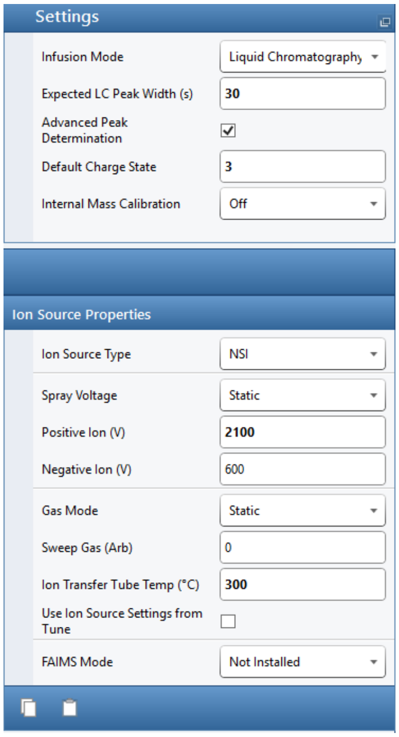

7. Click on **Scan Parameters**, and set to **Standard View.**

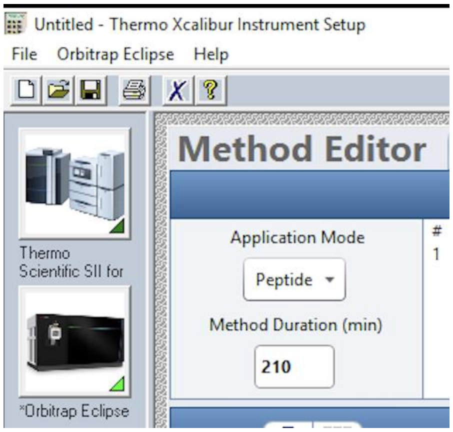

8. On the **Scans** menu, click **MS** and drag **Full Scan** into the **Experiment #1** window where it says **Place Scan Here** to set **Experiment #1** to **MS OT**.

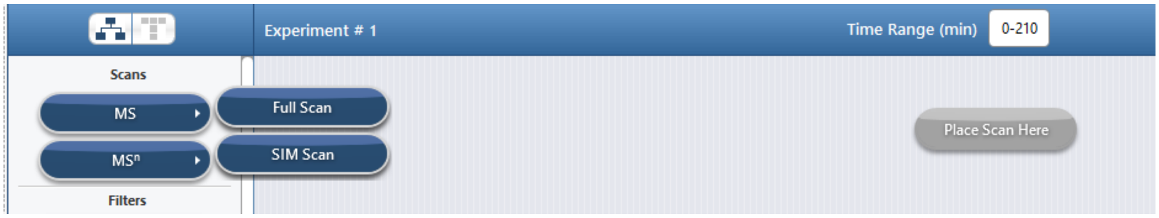

9. On the **MS Scan Properties** menu, change the following parameters:

- set the **Orbitrap Resolution** to 120000
- set the **Scan Range (m/z)** to 350 – 1650
- set the **AGC Target** to Custom
- set the **Normalized AGC Target (%)** to 750
- set the **Maximum Injection Time Mode** to Custom
- set the **Maximum Injection Time (ms)** to 60.

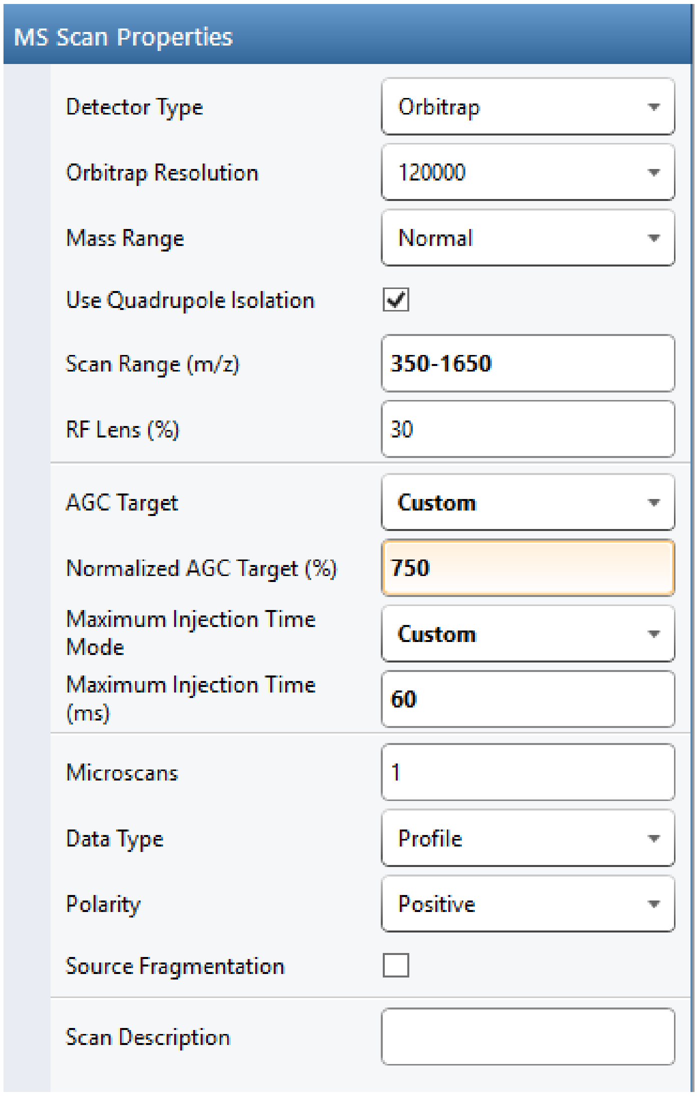

10. Under the **Experiment Actions** dropdown menu, click **Add New Experiment**.

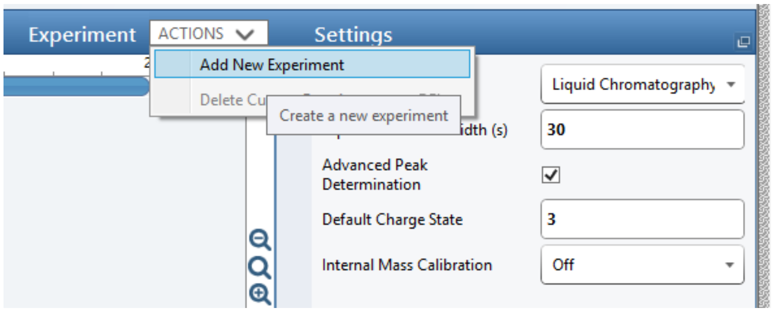

11. On the **Scans** menu, click **MS$^n$** and drag **MS$^n$** into the **Experiment #2** window where it says **Place Scan Here** to set **Experiment #2** to **tMS$^2$ OT CID.**
12. Under the **Targeted MS$^n$ Scan Properties** menu, change the following parameters:
    - set the **MS$^n$ Level (n)** to 2
    - set the **Isolation Mode** to Quadrupole
    - set the **Isolation Window (m/z)** to 1.6
    - set the **Activation Type** to HCD
    - set the **Collision Energy Mode** to Fixed
    - set the **HCD Collision Energy (%)** to 27
    - set the **Detector** to Orbitrap
    - set the **Orbitrap Resolution** to 30000
    - set **TurboTMT** to Off
    - set the **Mass Range** to Normal
    - set the **Scan Range Mode** to Define First Mass
    - set the **First Mass (m/z)** to 200
    - set the **RF Lens (%)** to 30
    - set the **AGC Target** to Custom
    - set the **Normalized AGC Target (%)** to defined in table by clicking the icon

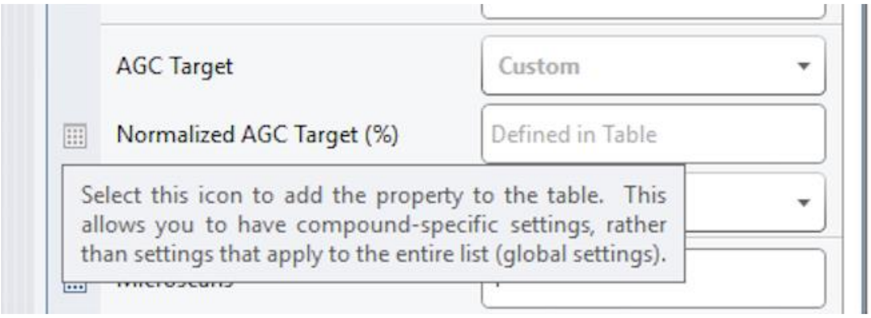

- set the **Maximum Injection Time Mode** to Auto
- set the number of **Microscans** to 1
- set the **Data Type** to Profile
- set the **Polarity to** Positive
- set the **Loop Control** to N
- set the **N (Number of Spectra)** to 26
- set the **Dynamic Retention Time** to Off
- set the **Time Mode** to Start/End Time.

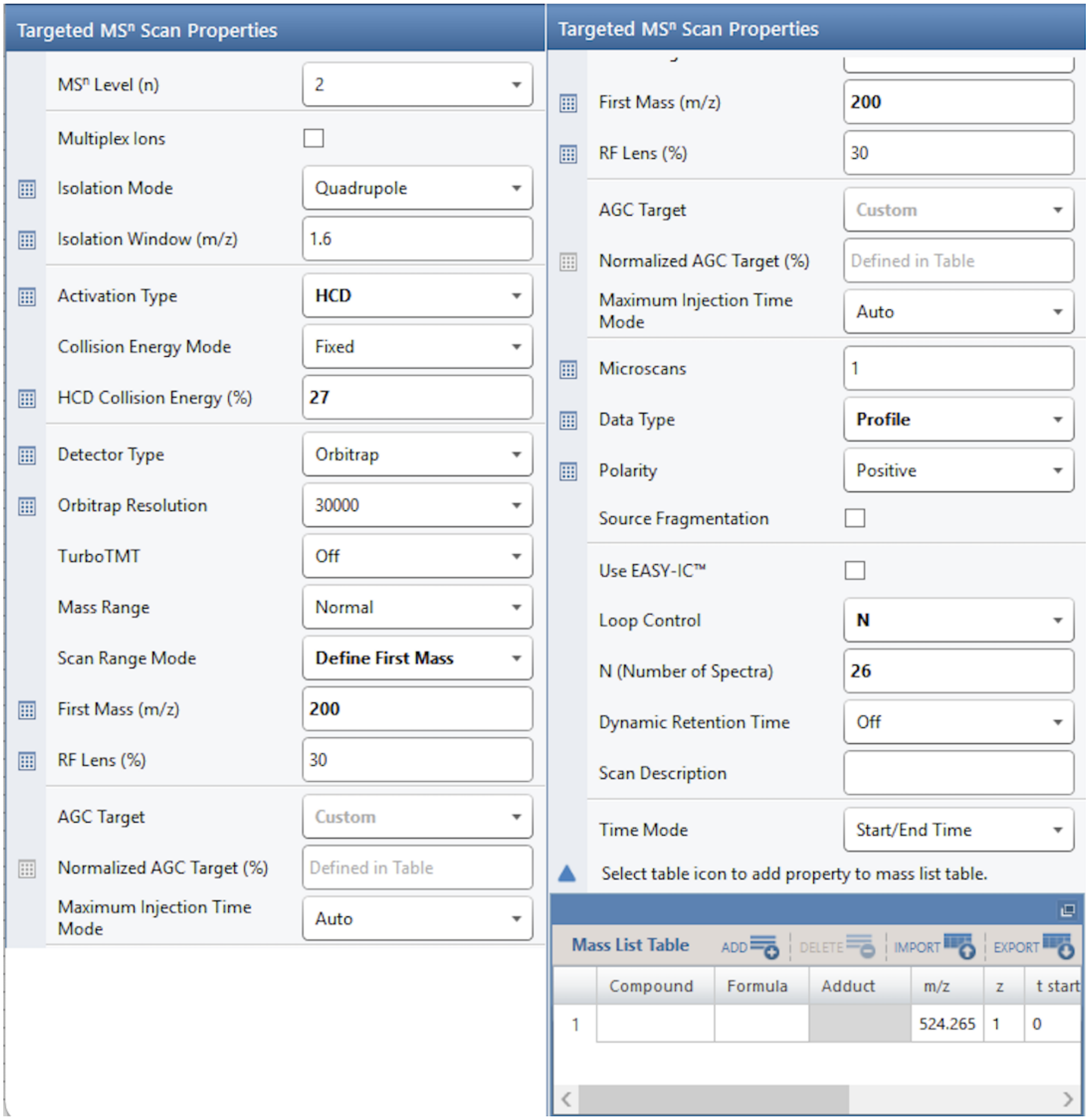

13. Import the isolation window strategy to the **Mass List Table** in the **Targeted MS$^n$ Scan Properties** menu:

- prepare the **DIA Variable Window text file** from the Supplementary Table S1 that defines the variable width isolation window strategy
- in the **Mass List Table** click **Import**
- select the **DIA Variable Window text file**
- ensure that **t start (min)** and **t stop (min)** in the **Mass List Table** corresponds to the length of the gradient.

Mass List Table

ADD DELETE IMPORT EXPORT

| | Compound | Formula | Adduct | m/z | z | t start (min) | t stop (min) | Isolation Window (m/z) | Normalized AGC Target (%) |
|---|---|---|---|---|---|---|---|---|---|
| 1 | | | | 366.5 | 3 | 0 | 210 | 33 | 6000 |
| 2 | | | | 395 | 3 | 0 | 210 | 26 | 6000 |
| 3 | | | | 418 | 3 | 0 | 210 | 22 | 6000 |
| 4 | | | | 438 | 3 | 0 | 210 | 20 | 6000 |
| 5 | | | | 457 | 3 | 0 | 210 | 20 | 6000 |
| 6 | | | | 475 | 3 | 0 | 210 | 18 | 6000 |
| 7 | | | | 493 | 3 | 0 | 210 | 20 | 6000 |
| 8 | | | | 511.5 | 3 | 0 | 210 | 19 | 6000 |
| 9 | | | | 529.5 | 3 | 0 | 210 | 19 | 6000 |
| 10 | | | | 547.5 | 3 | 0 | 210 | 19 | 6000 |
| 11 | | | | 565.5 | 3 | 0 | 210 | 19 | 6000 |
| 12 | | | | 584 | 3 | 0 | 210 | 20 | 6000 |
| 13 | | | | 603.5 | 3 | 0 | 210 | 21 | 6000 |
| 14 | | | | 623.5 | 3 | 0 | 210 | 21 | 6000 |
| 15 | | | | 644.5 | 3 | 0 | 210 | 23 | 6000 |
| 16 | | | | 666.5 | 3 | 0 | 210 | 23 | 6000 |
| 17 | | | | 689 | 3 | 0 | 210 | 24 | 6000 |
| 18 | | | | 713 | 3 | 0 | 210 | 26 | 6000 |
| 19 | | | | 740.5 | 3 | 0 | 210 | 31 | 6000 |
| 20 | | | | 771 | 3 | 0 | 210 | 32 | 6000 |
| 21 | | | | 804.5 | 3 | 0 | 210 | 37 | 6000 |
| 22 | | | | 842 | 3 | 0 | 210 | 40 | 6000 |
| 23 | | | | 887.5 | 3 | 0 | 210 | 53 | 6000 |
| 24 | | | | 946 | 3 | 0 | 210 | 66 | 6000 |
| 25 | | | | 1027.5 | 3 | 0 | 210 | 99 | 6000 |
| 26 | | | | 1363 | 3 | 0 | 210 | 574 | 6000 |

14. Click **File** and **Save** the instrument method file.

#### Chapter 2. Data Pre-Processing (Optional Step)

Once the DIA data is acquired, there are several data processing pipelines available to process the data. As an example, a DIA data set was acquired using the above method, then pre-processed using the HTRMS Converter program before directDIA analysis using Spectronaut (Biognosys, Schlieren, Switzerland). The HTRMS file conversion is optional and decreases the time to search the data using Spectronaut by converting vendor specific file formats into a **H**igh **T**ime **R**esolution **M**ass **S**pectrometry (HTRMS) file format. The optional file conversion step is optimal for large sample sizes.

1. Install the HTRMS Convertor program from Biognosys onto the MS computer.

   *Note: This requires a restart.*

2. Open the **HTRMS Convertor program** to set the program up to convert acquired data-independent acquisition files to a *.htrms peaks list in a monitored folder.
3. Select **Add Folder**.

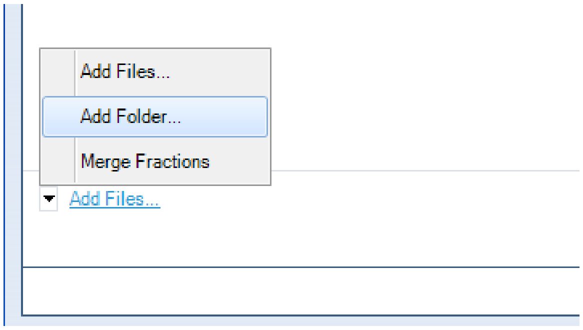

4. Select the **Source** and **Destination Directories**.
5. Ensure that **Monitor Folder** is enabled.
6. Click **Ok**.

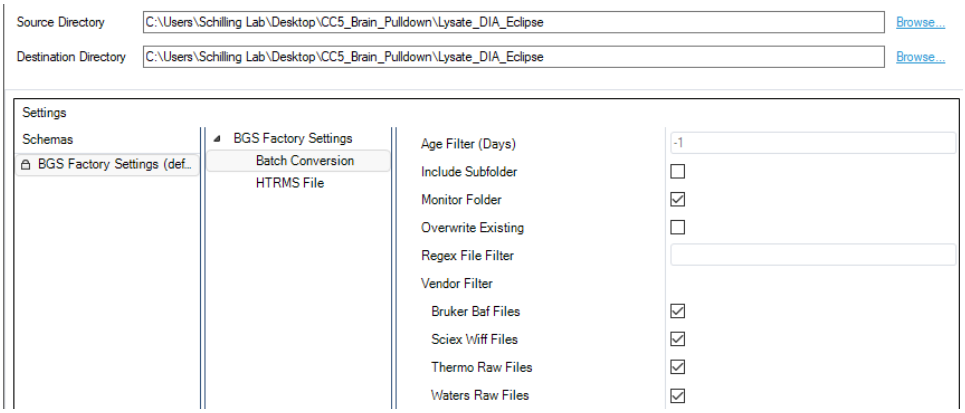

### Chapter 3. Data Processing

Spectronaut data processing using directDIA analysis searches each sample acquisition to form a spectral library, then searches each sample acquisition against the library to identify precursors, peptides, proteins, and protein groups.

1. Open **Spectronaut.**

   *Note: This tutorial is based on Spectronaut version 15.7.220308.50606.*

2. Select the **Pipeline** tab:
   - click **Set up a directDIA analysis from File**
   - navigate to the experimental vendor specific files or *.htrms files and select all files pertinent to the experiment
   - Click **Open**.

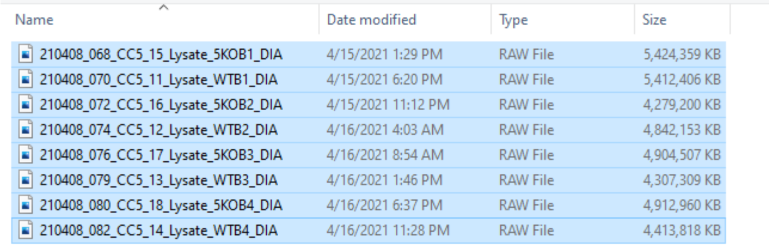

3. Name the experiment with a unique identifier.

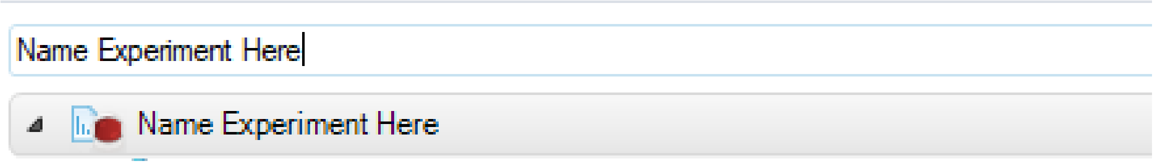

4. Click **Next**.
5. Select the appropriate organism **protein database**.

   *Note: In this study, the FASTA file "uniprot_mouse_proteome" is used.*

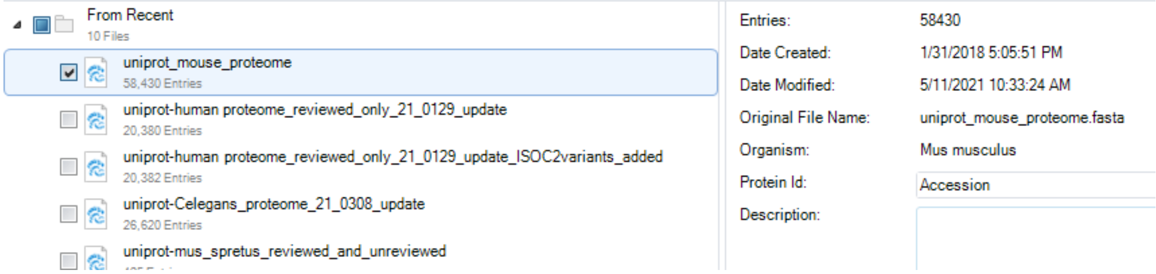

6. Click **Next**.

### Chapter 3.1. Protein Lysate: directDIA Search Settings

Create a **new search and extraction settings schema** for directDIA analysis of a digested protein lysate without specific post-translational modifications. Defaults should be preserved unless specifically mentioned in the text below.

1. Navigate to the **Pulsar Search Peptides** tab:

- the *Enzymes/Cleavage Rules* should be set to “Trypsin/P”
- set the *Digest Type* to “Specific”
- set the *Missed Cleavages* to “2”
- enable *Toggle N-terminal M.*

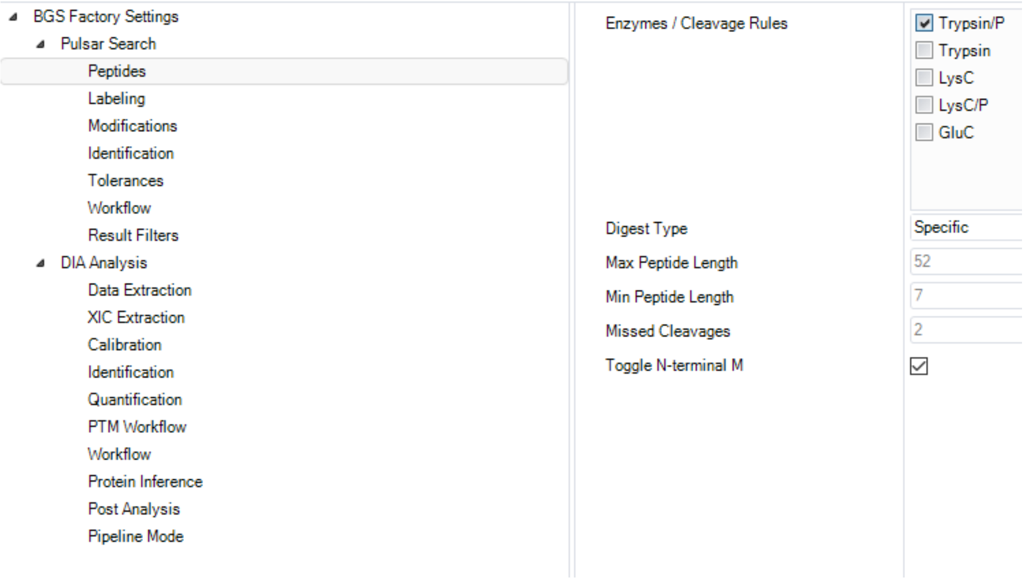

2. Select the **Pulsar Search Modifications** tab:

- set the *Max Variable Modifications* to “5”
- set the *Fixed Modifications* to “Carbamidomethyl (C)”
- set the *Variable Modifications* to “Acetyl (Protein *N*-Term)” and “Oxidation (M)”.

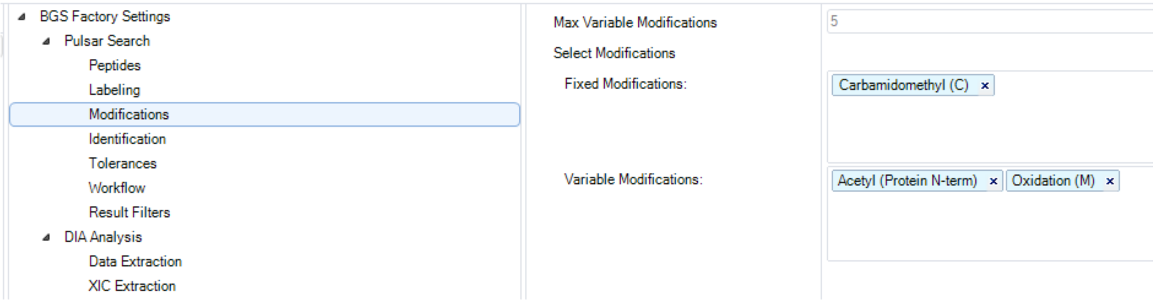

3. Navigate to the **DIA Analysis Identification** tab:

- set the *Precursor PEP Cutoff* to “1”
- set the *Precursor Qvalue Cutoff* to “0.01”
- set the *Protein Qvalue Cutoff (Experiment)* to “0.01”
- set the *Protein Qvalue Cutoff (Run)* to “0.05”.

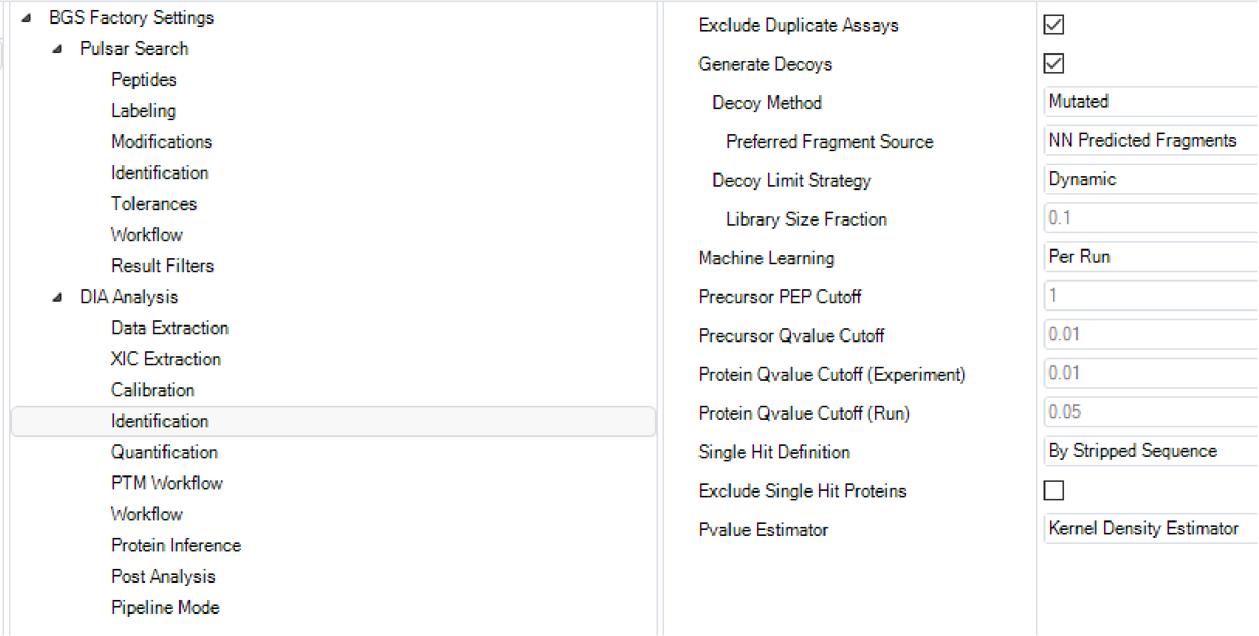

4. Click on the **DIA Analysis Quantification** tab:

- set the *Minor (Peptide) Grouping* to “by Stripped Sequence”
- set the *Major Group Quantity to “Sum peptide quantity”*
- set the *Major Group Max* to “7”
- set the *Major Group Min* to “1”
- set *Minor Group Quantity* to “Sum precursor quantity”
- set the *Minor Group Max* to “10”
- set the *Minor Group Min* to “1”
- set the *Data Filtering* to “Qvalue sparse”
- set the *Imputing Strategy* to “No Imputing”
- set the *Normalization Strategy* to “Local Normalization”
- set the *Row Selection* to “Q-value sparse”.

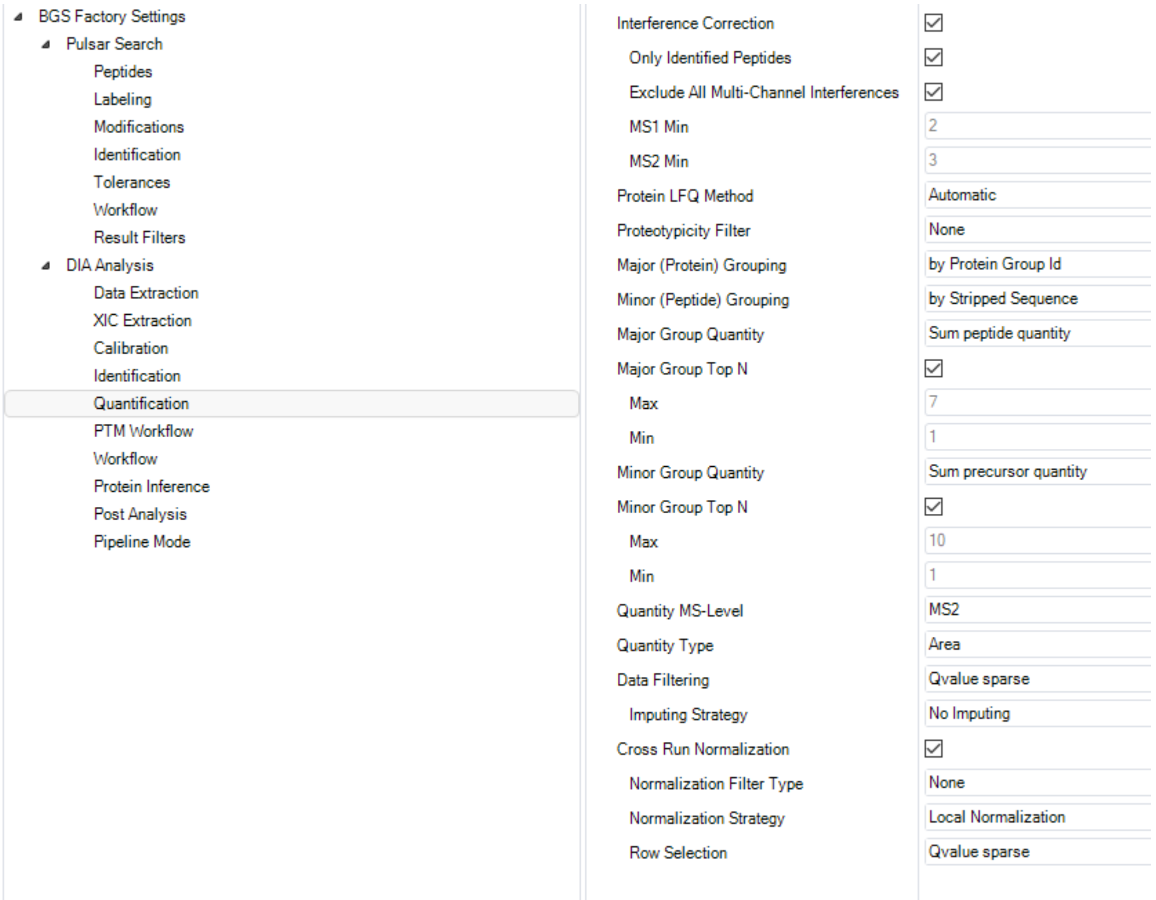

5. Select the **DIA Analysis Workflow** tab”

- set the *Profiling Strategy* to “iRT Profiling”.

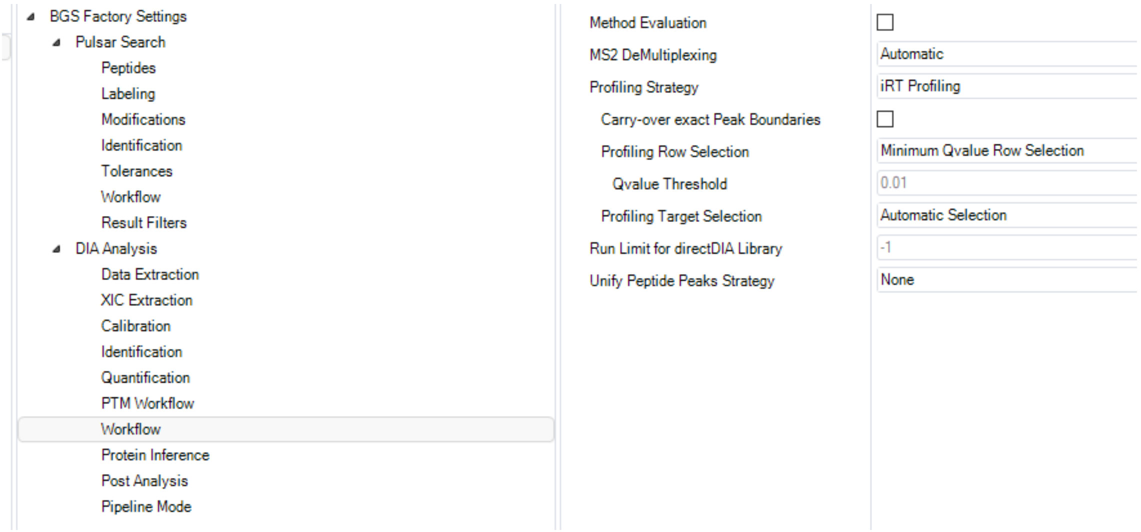

6. Navigate to the **DIA Analysis Post Analysis** tab:

- set the *Smallest Quantitative Unit* to “Precursor Ion (Quantification Settings)”
- set the *Differential Abundance Testing* to “Paired t-test”
- enable *Group-Wise Testing Correction*.

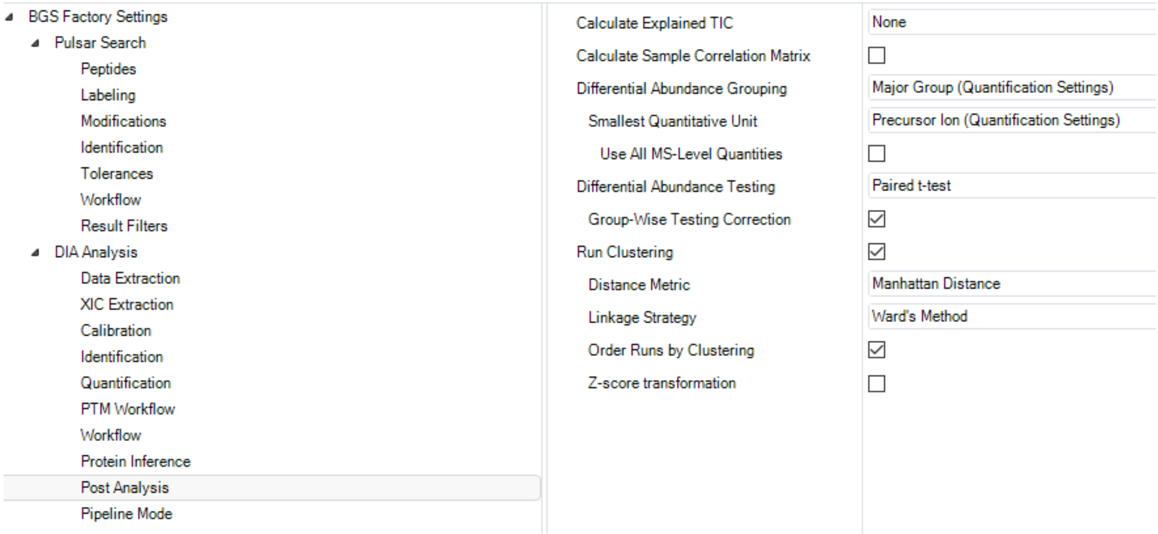

7. Click on the **DIA Analysis Pipeline Mode** tab:

- enable *Generate SNE File*
- select the appropriate *Report Schema* as described in **Chapter 4. Data Reporting***:* “Protein_Quant_Pivot” and “Peptide_Quant_Pivot”.

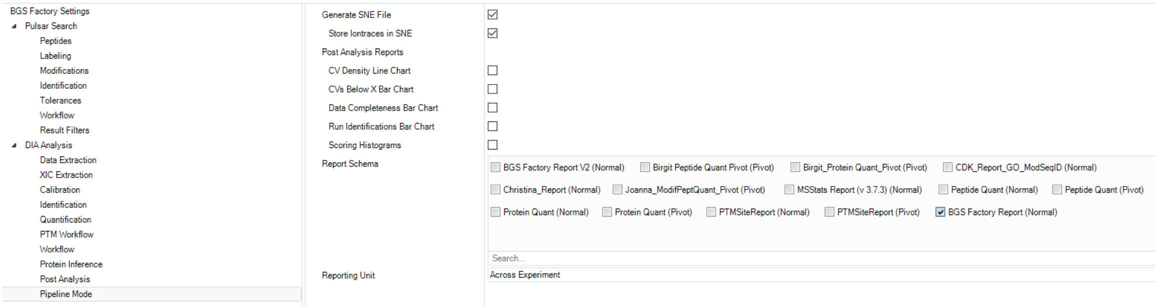

8. Click **Next**.

9. Go to **Chapter 3.3** to complete modifying search settings.

### Chapter 3.2. Post-translational Modifications: directDIA Search Settings

Create a **new search and extractions settings schema** for directDIA of a digested protein lysate enriched for specific post-translational modified peptides.

*Note: Below is an example of a schema for directDIA analysis of a digested protein lysate enriched for succinylated peptides.*

1. Navigate to the **Pulsar Search Peptides** tab

- set the *Enzymes/Cleavage Rules* to "Trypsin/P"
- set the *Digest Type* to "Specific"
- set *Missed Cleavages* to "2"
- enable *Toggle N-terminal M*.

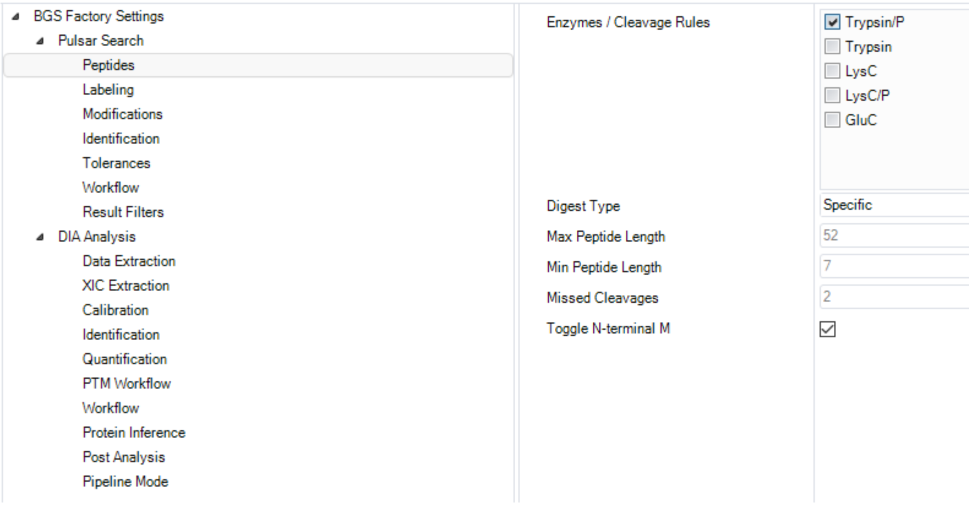

2. Select the **Pulsar Search Modifications** tab:

- set the *Max Variable Modifications* to "8"
- set the *Fixed Modifications* to "Carbamidomethyl (C)"
- set the *Variable Modifications* to "Acetyl (Protein *N*-Term)", "Oxidation (M)", and "Succinyl".

  *Note: Update "Succinyl" with the post-translational modification(s) of interest.*

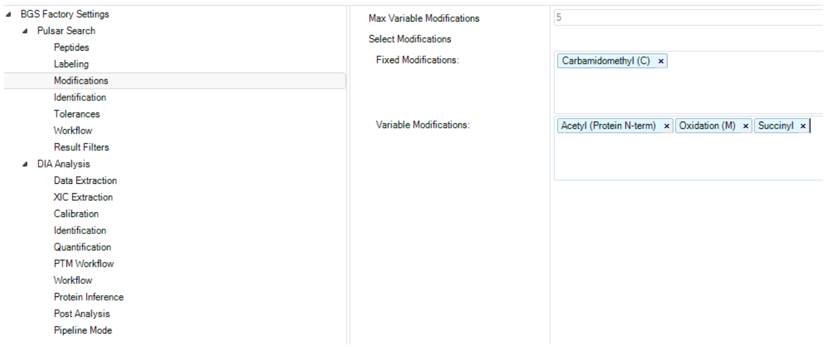

3. Navigate to the **DIA Analysis Identification** tab:

- set the *Precursor PEP Cutoff* to “1”
- set the *Precursor Qvalue Cutoff* to “0.01”
- set the *Protein Qvalue Cutoff (Experiment)* to “0.01”
- set the *Protein Qvalue Cutoff (Run)* to “1”.

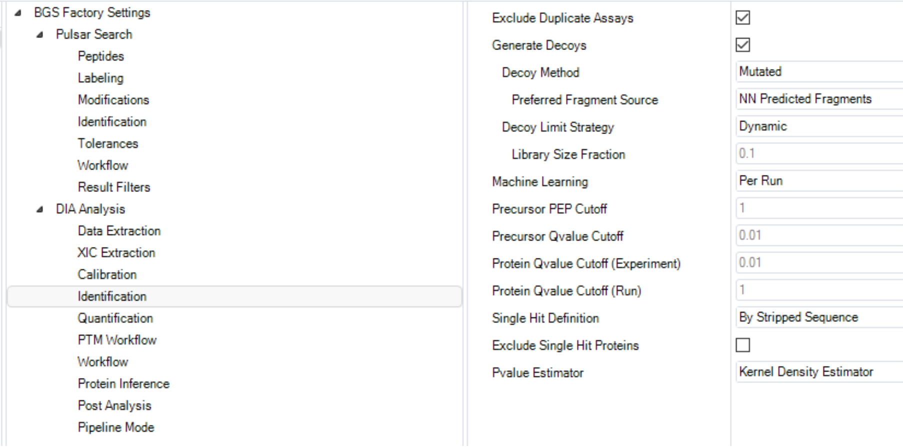

4. Select the **DIA Analysis Quantification** tab:

- set the *Minor (Peptide) Grouping* to “Modified sequence”
- set the *Major Group Quantity to “*Mean peptide quantity*”*
- set the *Major Group Max* to “3” and Min to “1”
- set *Minor Group Quantity* to “Mean precursor quantity”
- set the *Minor Group Max* to “3” and Min to “1”
- set the *Data Filtering* to “Qvalue sparse”
- set the *Imputing Strategy* to “No Imputing”
- ensure that the *Normalization Strategy* is unchecked.

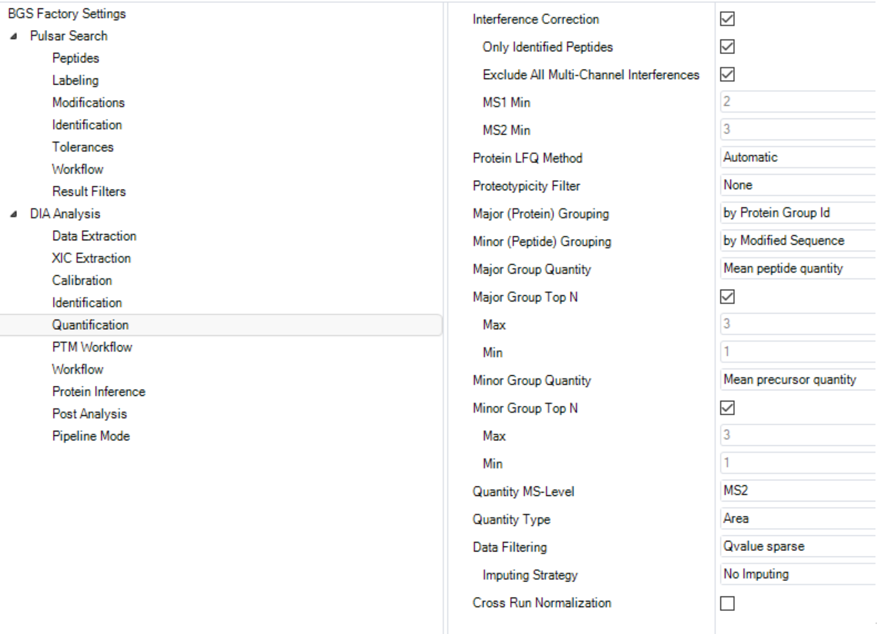

5. Click on the **DIA Analysis PTM Workflow** tab:

- enable *PTM Localization*
- set the *Probability Cutoff* to “0.75”.

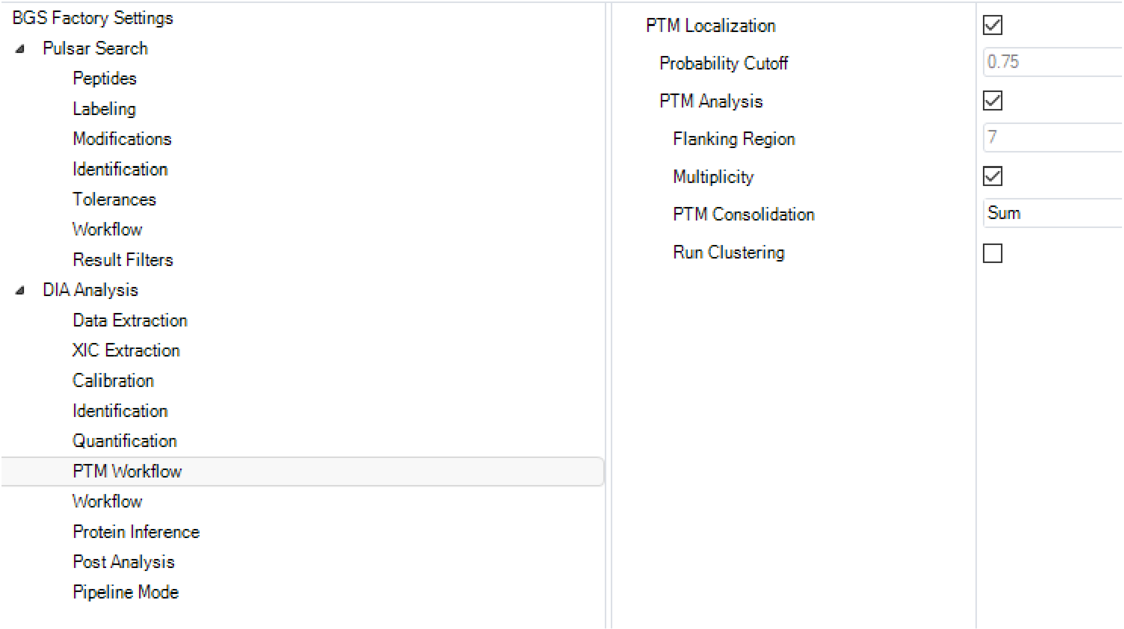

6. Select the **DIA Analysis Workflow** tab

- set the *Profiling Strategy* to “None”.

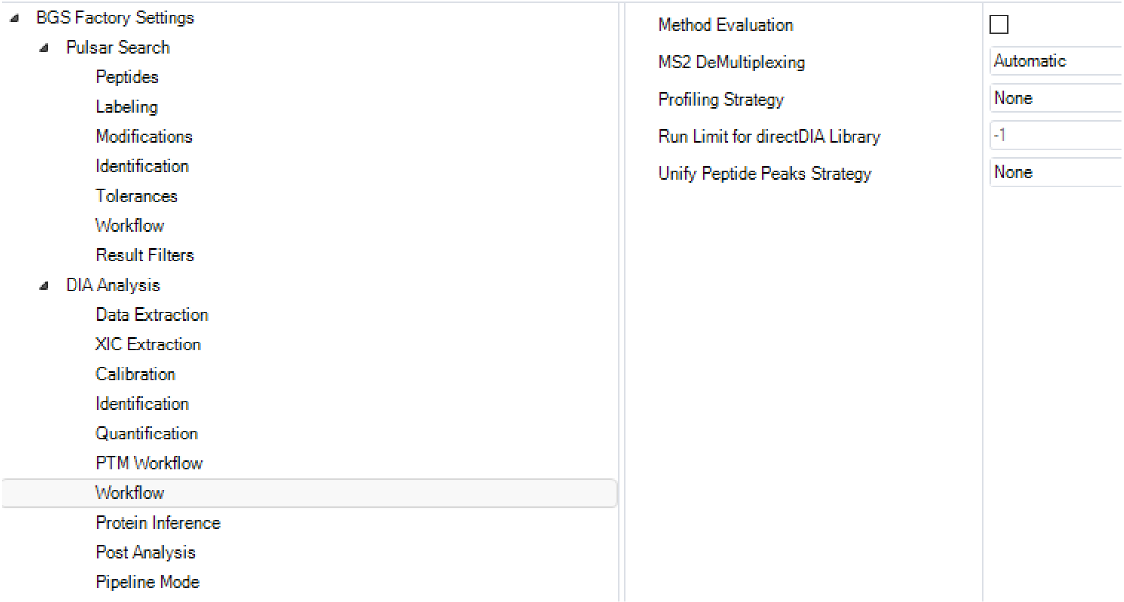

7. Navigate to the **DIA Analysis Post Analysis** tab:

- set the *Differential Abundance* Grouping to “Minor Group (Quantification Settings)”
- the *Smallest Quantitative Unit* should be set to “Precursor Ion (Quantification Settings)”
- set *Differential Abundance Testing* to “Paired t-test”
- ensure that *Group-Wise Testing Correction* is disabled.

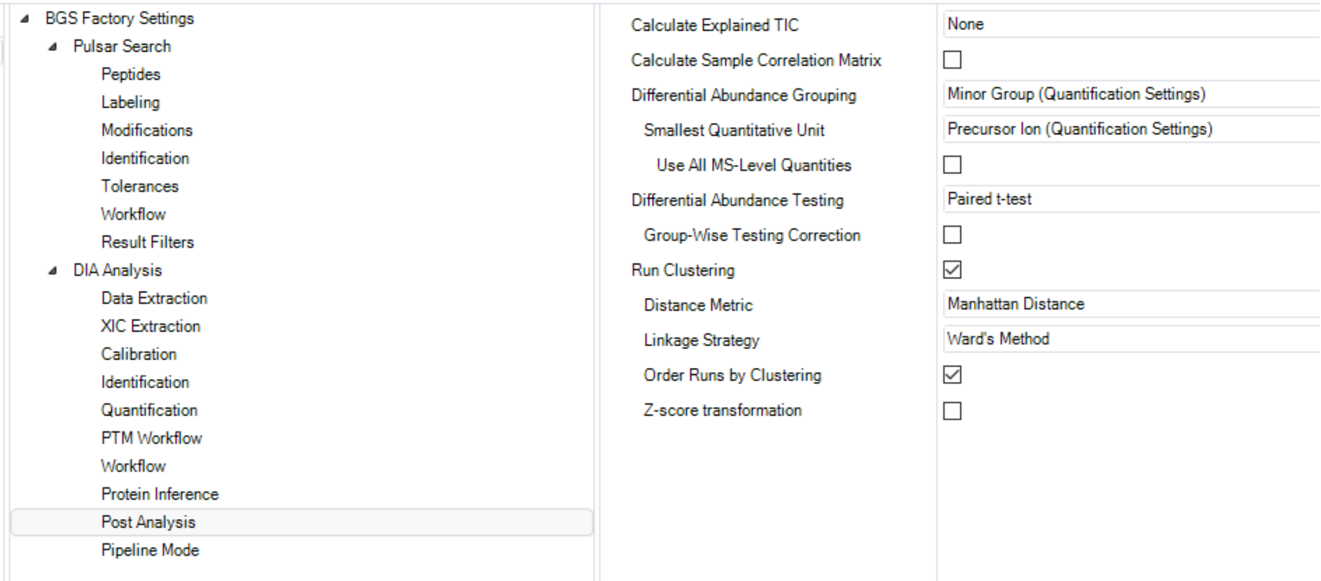

8. Click on the **DIA Analysis Pipeline Mode** tab:

- enable *Generate SNE File*
- select the appropriate *Report Schema* as described in **Chapter 4. Data Reporting***:* “Peptide_Quant_Pivot” and “PTM_Localization”.

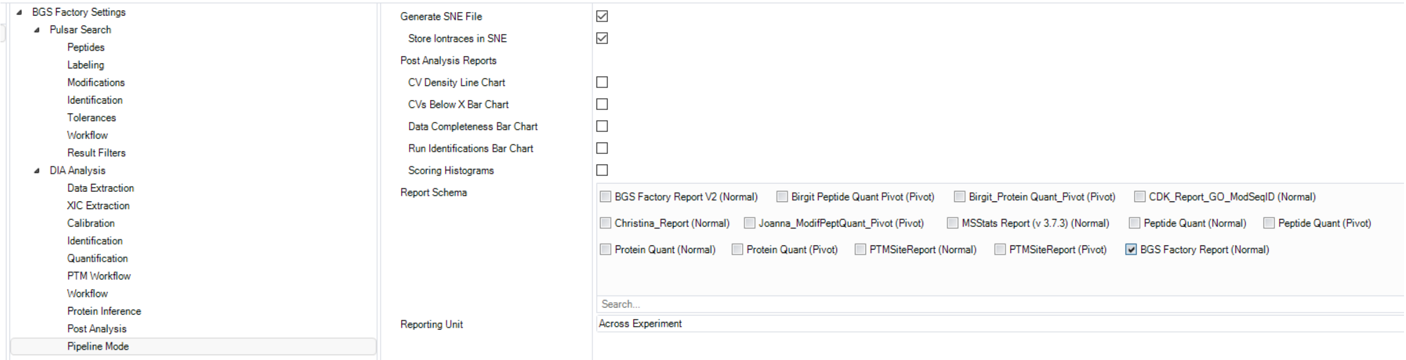

9. Go to **Chapter 3.3** to complete modifying search settings.

#### Chapter 3.3. directDIA Search Settings for Protein Lysate and Post Translational Modifications

1. Set the conditions for each experimental vendor specific file or *.htrms file included in the experiment.

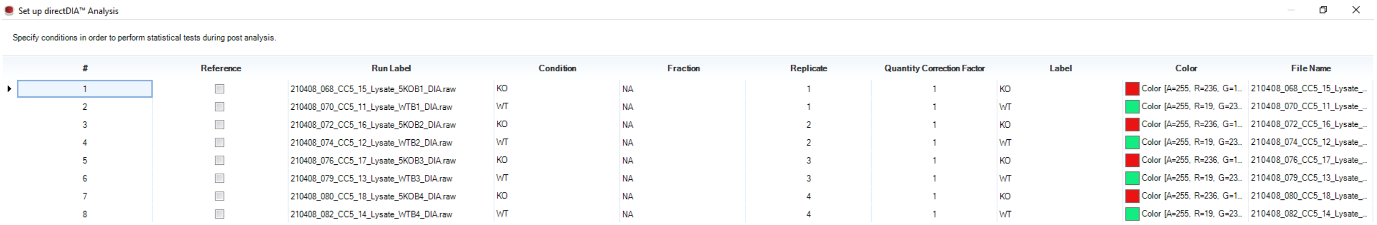

| # | Reference | Run Label | Condition | Fraction | Replicate | Quantity Correction Factor | Label | Color | File Name |
|---|---|---|---|---|---|---|---|---|---|
| 1 | ☐ | 210408_068_CC5_15_Lysate_5KOB1_DIA.raw | KO | NA | 1 | 1 | KO | Color [A=255, R=236, G=1... | 210408_068_CC5_15_Lysate_... |
| 2 | ☐ | 210408_070_CC5_11_Lysate_WTB1_DIA.raw | WT | NA | 1 | 1 | WT | Color [A=255, R=19, G=23... | 210408_070_CC5_11_Lysate_... |
| 3 | ☐ | 210408_072_CC5_16_Lysate_5KOB2_DIA.raw | KO | NA | 2 | 1 | KO | Color [A=255, R=236, G=1... | 210408_072_CC5_16_Lysate_... |
| 4 | ☐ | 210408_074_CC5_12_Lysate_WTB2_DIA.raw | WT | NA | 2 | 1 | WT | Color [A=255, R=19, G=23... | 210408_074_CC5_12_Lysate_... |
| 5 | ☐ | 210408_076_CC5_17_Lysate_5KOB3_DIA.raw | KO | NA | 3 | 1 | KO | Color [A=255, R=236, G=1... | 210408_076_CC5_17_Lysate_... |
| 6 | ☐ | 210408_079_CC5_13_Lysate_WTB3_DIA.raw | WT | NA | 3 | 1 | WT | Color [A=255, R=19, G=23... | 210408_079_CC5_13_Lysate_... |
| 7 | ☐ | 210408_080_CC5_18_Lysate_5KOB4_DIA.raw | KO | NA | 4 | 1 | KO | Color [A=255, R=236, G=1... | 210408_080_CC5_18_Lysate_... |
| 8 | ☐ | 210408_082_CC5_14_Lysate_WTB4_DIA.raw | WT | NA | 4 | 1 | WT | Color [A=255, R=19, G=23... | 210408_082_CC5_14_Lysate_... |

2. Click *Export Condition Setup*, name the file, and click **Save**.
3. Click **Next.**
4. Select the appropriate **Gene Ontology Annotations** for the experiment.

   *Note: In this study, the file* “Mus musculus (GO Annotations Uniprot)” *is used.*

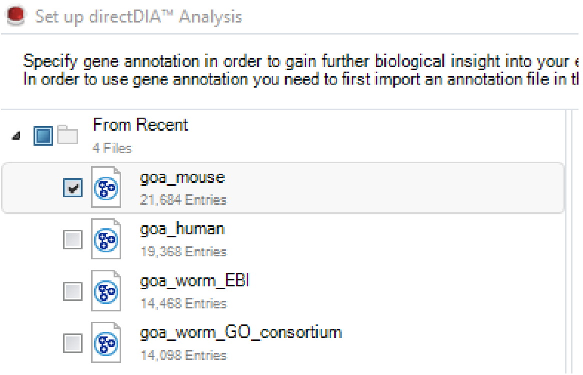

5. Click **Next**.
6. Click **Browse**:
   - select the appropriate **Output Directory** to save the sne. file and export the reports, plots, and candidate protein files.

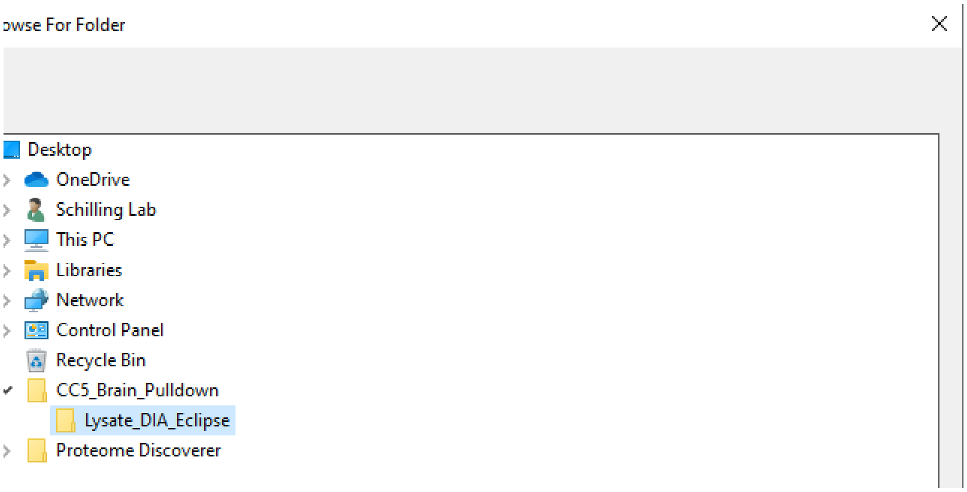

7. Click **Ok**.
8. Click **Finish** to queue the analysis.
9. Click **Run Pipeline** to start the analysis queue.

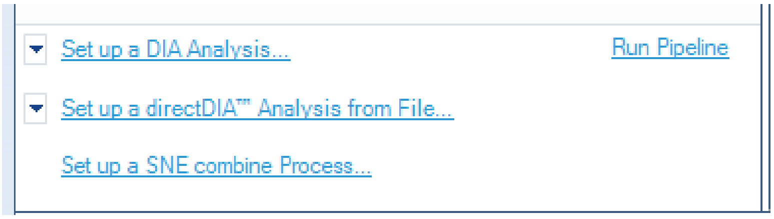

### Chapter 4. Data Reporting

#### Chapter 4.1. Protein Quantification Report

1. Navigate to the **Report** tab to set up a custom report.
2. Select the **BGS Factory Report** under the **Run Pivot Report** dropdown to create a custom Protein Quantification Pivot Report, “Protein_Quant_Pivot”.
3. Select the following **Row Labels**:
   - PG.Qvalue
   - PG.Genes
   - PG.ProteinDescriptions
   - PG.UniProtIDs
   - PG.ProteinNames
   - PG.CellularComponent
   - PG.BiologicalProcess
   - PG.MolecularFunction.

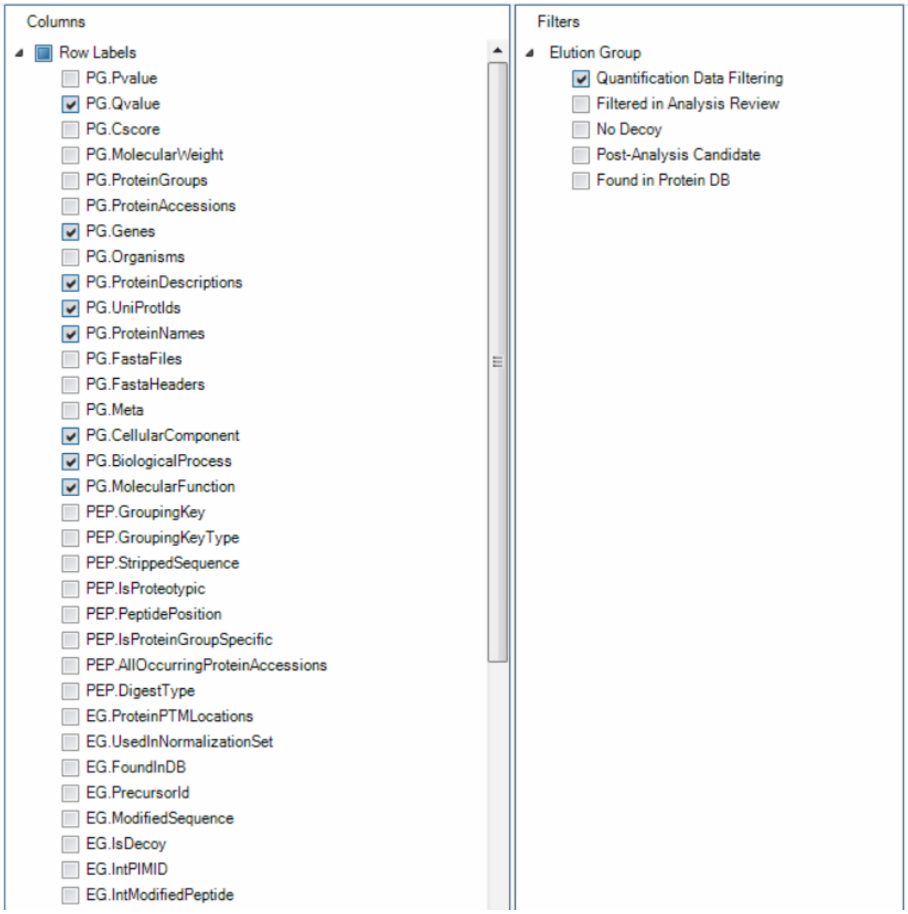

4. Select the following **Cell Values**:

- PG.NrOFPrecursorsIdentified
- PG.NrOFPrecursorsUsedForQuantification
- PG.Quantity.

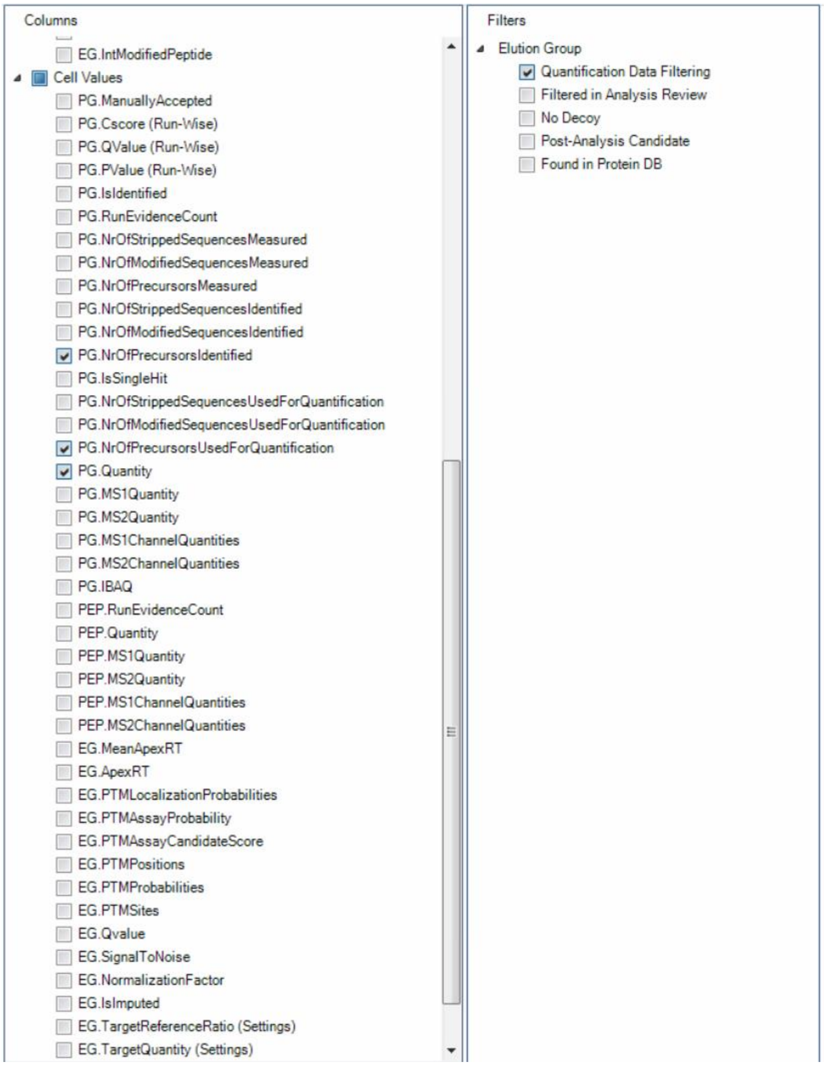

5. Select the following **Filters**:

- Quantification Data Filtering.

6. Select **Save As** and save the report scheme as “Protein_Quant_Pivot”.

#### Chapter 4.2. Peptide Quantification Report

1. Navigate to the **Report** tab to set up a custom report.
2. Select the **BGS Factory Report** under the **Run Pivot Report** dropdown to create a custom Peptide Quantification Pivot Report, “Peptide_Quant_Pivot”.
3. Select the following **Row Labels**:
   - PG.Qvalue
   - PG.Genes
   - PG.ProteinDescriptions
   - PG.UniProtIDs
   - PG.ProteinNames
   - PG.CellularComponent
   - PG.BiologicalProcess
   - PG.MolecularFunction
   - PEP.PeptidePosition
   - EG.PrecursorId
   - EG.ModifiedSequence.

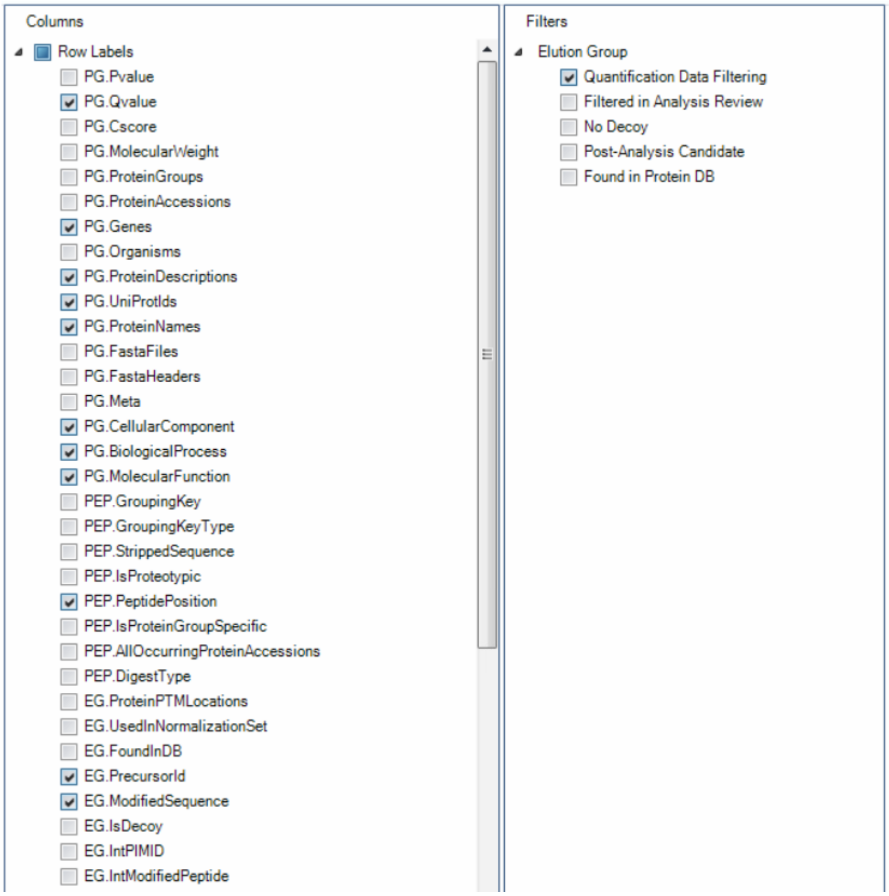

4. Select the following **Cell Values**:

- PEP.MS2Quantity
- EG.PTMProbabilities
- EG.PTMSites
- EG.TotalQuantity (Settings).

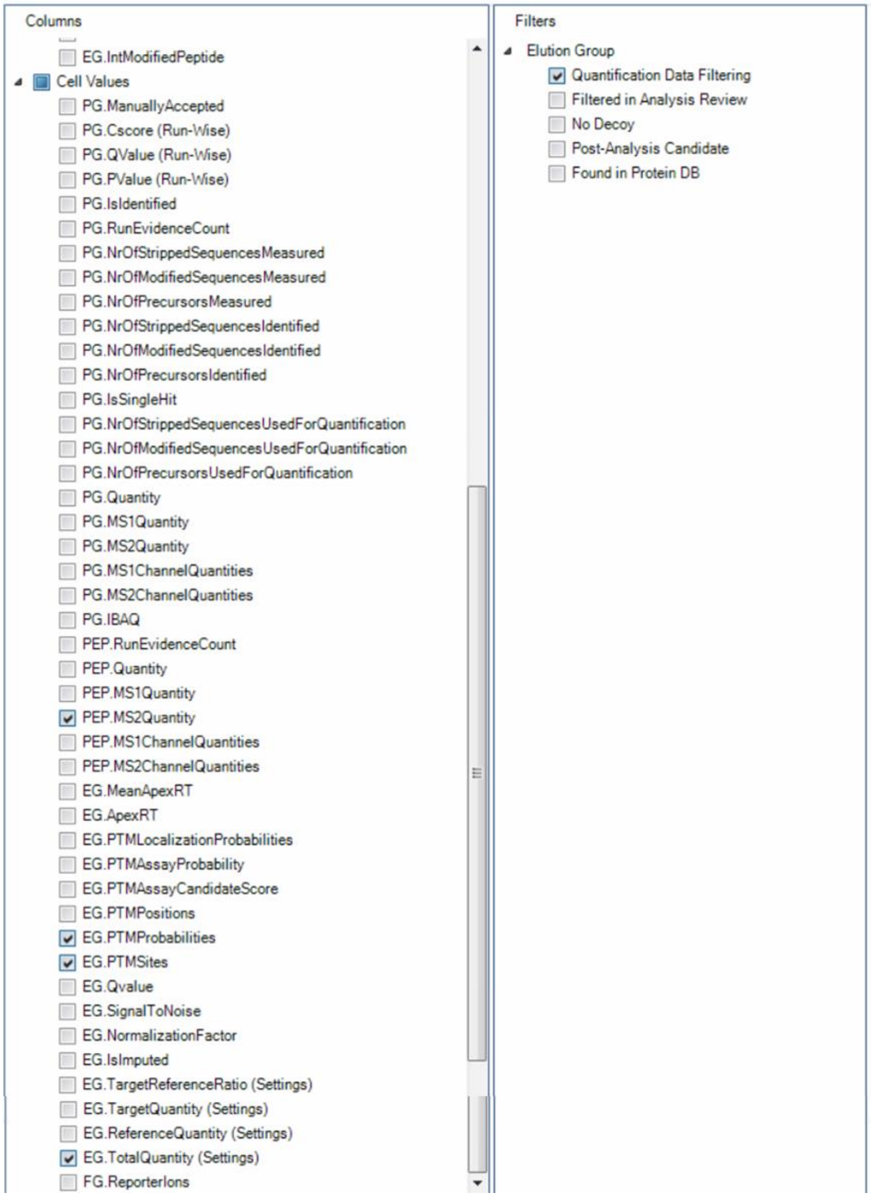

5. Select the following **Filters**:

- Quantification Data Filtering

6. Select **Save As** and save the report scheme as “Peptide_Quant_Pivot”.

#### Chapter 4.3. PTM Site Localization Report

1. Navigate to the **Report** tab to set up a custom report.
2. Select the **BGS Factory Report (default)** under the **PTM Site Report** tab and **Normal Report** dropdown to create a custom PTM Site Localization Report, “PTM_Localization”.
3. Select the following **Run** settings:

- R.Condition
- R.FileName
- R.Fraction
- R.Label
- R.Replicate.

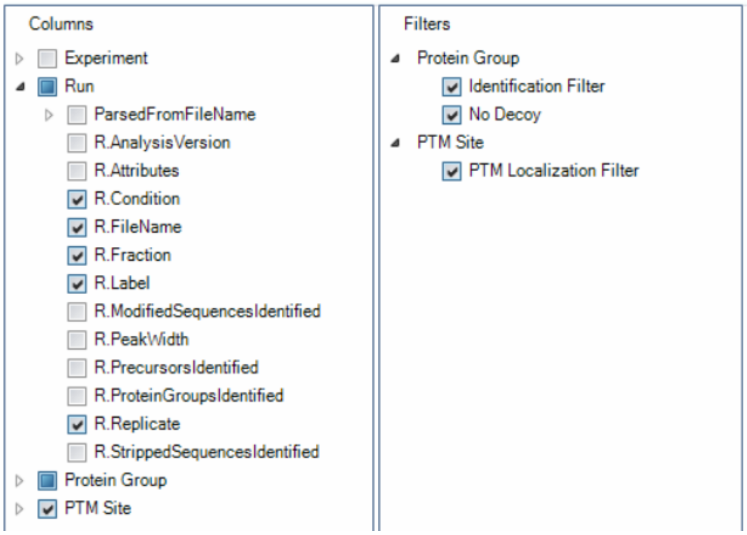

4. Select the following **Protein Group Assay** settings:

- PG.ProteinAccessions
- PG.ProteinGroups.

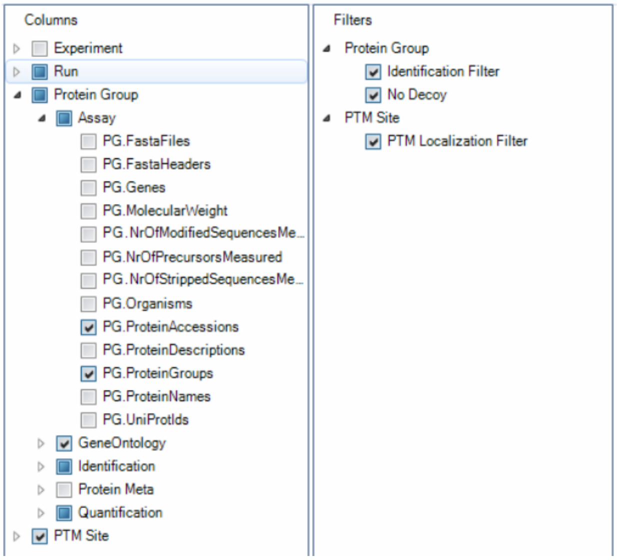

5. Select the following **Protein Group Gene Ontology** settings:

- PG.BiologicalProcess
- PG.CellularComponent
- PG.MolecularFunction.

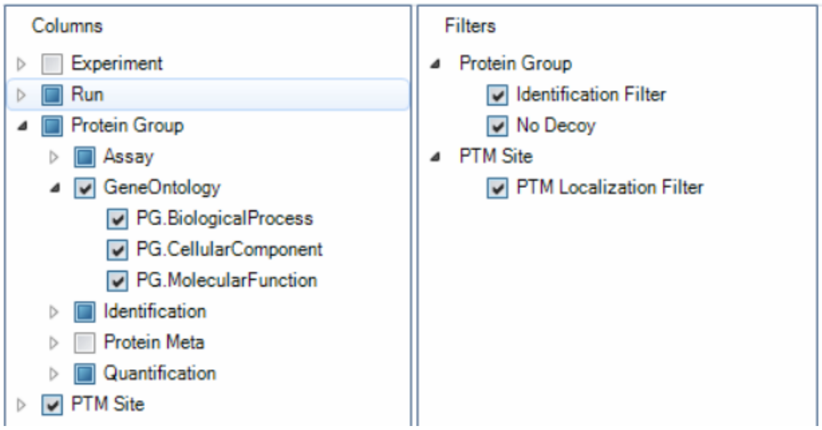

6. Select the following under **Protein Group Identification** settings:

- PG.Pvalue
- PG.Qvalue
- PG.Qvalue (Run-Wise).

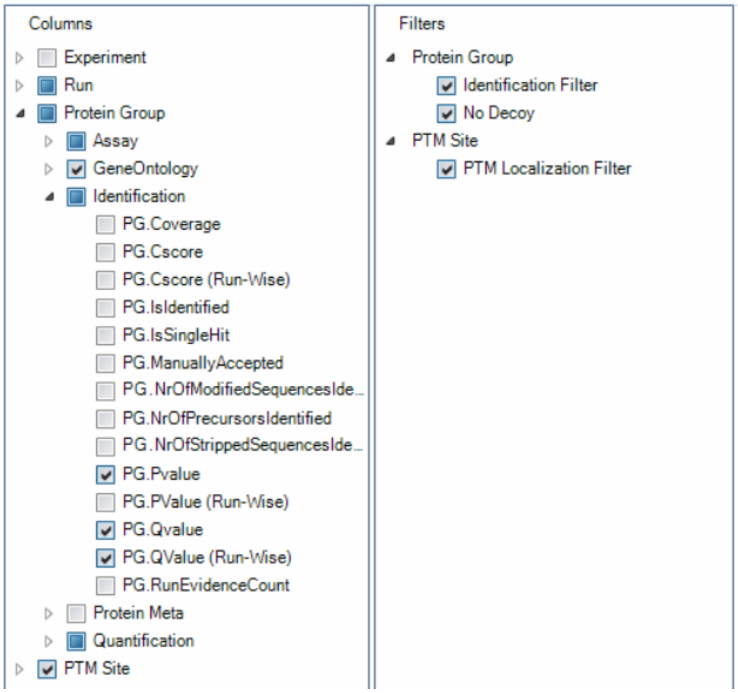

7. Select the following under **Protein Group Quantification** settings:

- PG.Quantity.

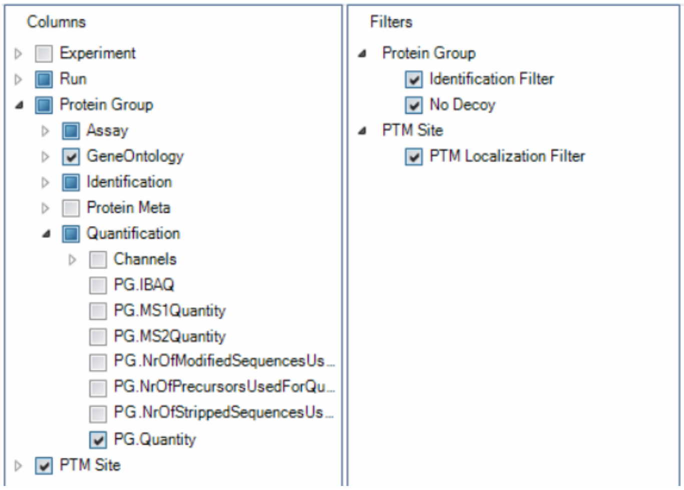

8. Select the following under **PTM Site** settings:

- PTM.CollapseKey
- PTM.FlankingRegion
- PTM.Group
- PTM.ModificationTitlte
- PT.Multiplicity
- PTM.NrOFCollapsedPeptides
- PTM.ProtienId
- PTM.Quantity
- PTM.SiteAA
- PTM.SiteLocalization
- PTM.SiteProbability

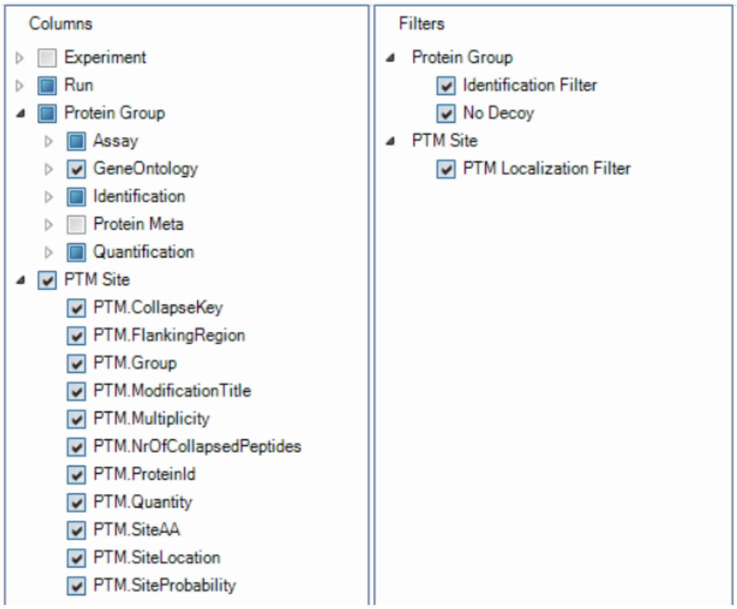

9. Select the following **Filters** settings:

- Identification Filter
- No Decoys
- PTM Localization Filter.

10. Select **Save As** and save the report scheme as “PTM_Localization”.
